# Network properties constrain natural selection on gene expression in *Caenorhabditis elegans*

**DOI:** 10.1101/2025.02.19.639144

**Authors:** Tyler R. Inskeep, Simon C. Groen

## Abstract

Gene regulatory networks (GRNs) integrate genetic and environmental signals to coordinate complex phenotypes and evolve through a balance of selection and drift. Using publicly available datasets from *Caenorhabditis elegans*, we investigated the extent of natural selection on transcript abundance by linking population-scale variation in gene expression to fecundity, a key fitness component. While the expression of most genes covaried only weakly with fitness, which is typical for polygenic traits, we identified seven transcripts under significant directional selection. These included *nhr-114* and *feh-1,* implicating variation in nutrient-sensing and metabolic pathways as impacting fitness. Stronger directional selection on tissue-specific and older genes highlighted the germline and nervous system as focal points of adaptive change. Network position further constrained selection on gene expression; high-connectivity genes faced stronger stabilizing and directional selection, highlighting GRN architecture as a key factor in microevolutionary dynamics. The activity of transcription factors such as *zip-3*, which regulates mitochondrial stress responses, emerged as targets of selection, revealing potential links between energy homeostasis and fitness. Our findings demonstrate how GRNs mediate the interplay between selection and drift, shaping microevolutionary trajectories of gene expression and phenotypic diversity.

## Introduction

Gene regulatory networks (GRNs) orchestrate organismal development and physiology and thus are critical to understanding phenotypic evolution. As populations diverge, changes in gene expression within GRNs can arise through both adaptive processes and neutral genetic drift, yet the relative contributions of these forces remain unresolved (True and Haag 2001; Hoekstra and Coyne 2007; Lynch 2007; Carroll 2008). Given that heritable variation in gene expression underlies variation for many complex traits (Wang *et al*. 1999; Carroll 2000; Ayroles *et al*. 2009; Zhang *et al*. 2022), deciphering how natural selection shapes GRNs is essential for understanding the mechanisms underlying phenotypic evolution (Romero *et al*. 2012).

GRN evolution is constrained by both network architecture and gene history. The “omnigenic model” posits that most gene expression differences in populations are mediated by genetic interactions with regulators in *trans* (Liu *et al*. 2019). Younger genes, often incorporated at the periphery of GRNs, tend to face relaxed selective constraints, facilitating their functional integration (Capra *et al*. 2010; Wei *et al*. 2016; Defoort *et al*. 2018). However, even well-conserved orthologs, such as those conserved among species in the nematode genus *Caenorhabditis,* exhibit weak coupling between rates of sequence and expression evolution, suggesting that network drift plays a significant role in the evolutionary dynamics of GRNs (True and Haag 2001; Castillo-Davis *et al*. 2004).

In multicellular organisms, functional interdependence between tissues constrains trait evolution (Liang *et al*. 2018). Tissue-specific GRNs can free genes from these constraints by minimizing pleiotropic consequences in different tissues. Tissue-specific genes often evolve more rapidly at the sequence level, while experiencing stabilizing selection on expression, reflecting the dual pressures of functional specialization and regulatory constraint (Zhang and Li 2004; Liao and Zhang 2006). Reproductive tissues stand out as hotspots for rapid GRN evolution, likely driven by sexual selection (Brawand *et al*. 2011; Fukushima and Pollock 2020; El Taher *et al*. 2021). Despite these insights, how GRN architecture and tissue specificity influence selection on gene expression at microevolutionary scales remains poorly understood.

Comparative studies of expression divergence have advanced our understanding of GRN evolution, yet they are often limited in their ability to disentangle adaptive evolution from lineage-specific drift (reviewed by Price *et al*. 2022). Population-level studies linking gene expression variation to fitness component metrics offer a complementary approach, shedding light on selective pressures in real time (Groen *et al*. 2020; Koch and Guillaume 2020; Ahmad *et al*. 2021). For instance, selection on population-scale variance in gene expression appears weak under certain conditions in rice (Groen *et al*. 2020) but biased towards diversifying selection in salmonid fish (Ahmad *et al*. 2021). These discrepancies contrast with the results of comparative approaches, which suggest that stabilizing selection dominates the transcriptome (Bedford and Hartl 2009; Kalinka *et al*. 2010; Chen *et al*. 2019; El Taher *et al*. 2021), raising critical questions about the modes of selection acting on GRNs across scales and species.

In this study, we leverage publicly available genome-wide gene expression and phenotypic data (Zhang *et al*. 2021, 2022) to investigate selection on transcript abundance in a population of field-collected strains of the nematode *Caenorhabditis elegans*, which have been reared in a laboratory environment for only a limited number of generations. Using genotypic selection analyses (Rausher 1992), we link gene expression variation to lifetime fecundity in this relatively new environment. To explore how GRN architecture influences microevolutionary rates of expression divergence, we integrate genome-wide regulatory data with information on gene age and tissue specificity. Finally, we use our models to look for evidence of reproduction-related tradeoffs specific to predominantly selfing species such as *C. elegans* (Cutter 2004; Murray and Cutter 2011). By examining these dynamics, our study provides new insights into the selective pressures shaping GRNs and the role of network architecture in their microevolutionary trajectories.

## Methods

### Univariate genotypic selection analyses

We estimated total (direct and indirect) linear (*S*) and quadratic (*C*) selection differentials for the abundance of 25,849 individual transcripts (representing approximately 16,094 genes) in a population of 207 genetically diverse strains of the *Caenorhabditis elegans* Natural Diversity Resource (CaeNDR)—as quantified by Zhang *et al*. (2022) from synchronized, laboratory-reared worms of each strain at the young adult stage—using the *glm()* function in R (Rausher 1992). To preprocess the genome-wide gene expression data for analysis, we excluded from the analysis expression information from 14 strains for which no total lifetime fecundity (TLF) data was available. As a fitness component metric, we normalized TLF measurements (normTLF) from worms reared in the same laboratory environment obtained by Zhang *et al*. (2021). We did this through dividing TLF of each strain by the population mean. After this, we standardized the abundance of each transcript (stand_expr) by subtracting the population mean for that transcript and dividing by the population standard deviation. For linear selection (*S*) models, we provided *glm()* with the formula ‘*normTLF ∼ stand_expr’,* and for quadratic selection *(C)*, we provided the formula ‘*normTLF ∼ stand_expr + I(stand_expr^2^)’*. Estimates of *C* were then multiplied by 2 (Rausher 1992). Bonferroni correction was performed to correct for multiple testing using the *p.adjust()* function in R.

To assess the impact of population structure on the strength and pattern of selection, information on divergent *C. elegans* strains was retrieved from Zhang *et al*. (2021). This revealed 21 strains as being highly divergent from the other strains in the population based on genome-wide diversity metrics. Selection differentials were then estimated using the same methods but with these 21 divergent strains excluded from the analysis. We additionally estimated total selection differentials for time to maturity, as predicted from the strain-specific transcriptomes in Zhang *et al*. (2022) following the same procedures.

### Multivariate genotypic selection analyses

We estimated direct linear (*β*) and quadratic (*Ɣ*) selection gradients on coordinated patterns of gene expression by performing principal component (PC) analysis using the *prcomp()* function in R. We selected the top 22 PCs that individually explained at least 0.5% of the population variance in genome-wide gene expression and retrieved loading values for the abundance of each transcript per strain into PC1-PC22. We submitted the strain-specific loading values to multivariate analysis using the *glm()* function in R with the formulas *’normTLF ∼ PC1 + PC2 + … + PC22’ (β)* and *’normTLF ∼ PC1 + I(PC1^2^) + PC2 + I(PC2^2^) + … + PC22 + I(PC22^2^)’ (Ɣ).* Estimates of *Ɣ* were then multiplied by 2 (Rausher 1992).

To obtain biological insight from PCs of interest, we retrieved the loading values for the abundance of each transcript into PC1-PC22. We considered the top 100 transcripts per PC for further analysis.

### Gene ontology (GO) analyses

To determine if selection acted on the expression of genes with unique functional categories, we retrieved Gene Ontology (GO) annotations for each gene from the Ensembl WormBase ParaSite BioMart release 19 (https://parasite.wormbase.org/biomart/martview) *C. elegans* (PRJNA13758, WS290) dataset. We merged our selection models with GO annotations and determined functional categories with distinct average selection strengths compared to the total genome-wide gene expression dataset using two-sided Mann-Whitney *U*-tests with the *wilcox.test()* function and adjusting the p-values using Bonferroni correction with the *p.adjust()* function in R.

Further gene set enrichment analyses (GSEAs) on smaller subsets of genes were performed using the ShinyGO v0.81 web server with all transcripts showing expression across the strains supplied as background (Ge *et al*. 2020).

### The influence of network architecture on the strength and pattern of selection

To determine how network position impacts the strength and pattern of selection on gene expression, we retrieved transcript-level expression quantitative trait loci (eQTL) determined by Zhang *et al*. (2022) using the same transcriptome data. We compared average selection differentials between transcripts that were or were not regulated by different types of eQTL using two-sided Mann-Whitney *U*-tests with the *wilcox.test()* function in R.

We retrieved information on accessible chromatin regions (ACRs) collected from bulk larval stage 4 (L4) worms in addition to ACRs from flow-sorted epidermal, hypodermal, intestinal, muscle, and germline nuclei (Serizay *et al*. 2020). ACRs, defined for 20,222 genes, were classified as promoters or enhancers based on genomic location (Serizay *et al*. 2020).

We also retrieved gene regulatory information from a consensus GRN built off of time-course RNA-sequencing data from aging worms (Suriyalaksh *et al*. 2022). The GRN included 313,562 interactions between 590 transcription factors (TFs) and 7,564 target genes. To enhance inference of TF-target relationships, the GRN was informed with gold-standard Chromatin Immunopurification (ChIP)-sequencing, yeast one hybrid assays, TF binding site analyses, and TF knockdown gene expression analyses (Suriyalaksh *et al*. 2022). We inferred in-degree as the number of TFs regulating a target and out-degree as the number of targets regulated by each TF.

To account for both the number of ACRs and GRN properties, we grouped genes into functional categories, ensuring that the size of each group was similar so that differences between the classes were minimized. We compared average selection differentials across the various regulatory categories using Kruskal-Wallis tests (*kruskal_test())* followed by Dunn’s post-hoc tests (*dunnTest()*) in R.

To estimate selection on TF activity, we determined groups of TF targets with significantly distinct average selection differentials relative to the set of 7,564 target genes using two-sided Mann-Whitney *U*-tests with the *wilcox.test()* function and adjusting the p-values using Bonferroni correction with the *p.adjust()* function in R. We determined the tissue of maximal expression using RNA-sequencing data from flow-sorted epidermal, hypodermal, intestinal, muscle, and germline nuclei described by Serizay *et al*. (2020).

### The influence of tissue specificity on the strength and pattern of selection

To determine how tissue specificity impacted the strength of selection on gene expression, we reanalyzed published transcript profiles of 20,222 genes across flow-sorted epidermal, hypodermal, intestinal, muscle, and germline nuclei (Serizay *et al*. 2020). For each gene, we determined the tissue specificity index *τ (tau):*

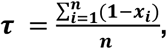

where *n* is the number of tissues profiled and *x_i_* is the normalized expression of a gene in a given tissue (Yanai *et al*. 2005). We grouped genes into ten equally-sized bins of tissue-specificity and compared average selection differentials across these bins using Kruskal-Wallis tests (*kruskal_test())* followed by Dunn’s post-hoc tests (*dunnTest()*) in R.

We retrieved information on genes expressed specifically in each profiled tissue: epidermis, hypodermis, intestine, muscle, germline, and sperm (Serizay *et al*. 2020). We then determined 95% confidence intervals for the distributions of *S* and *|S|* for tissue-specific gene sets and identified tissues facing stronger- or weaker-than average selection compared to the genome-wide confidence intervals.

### The influence of gene age on the strength and pattern of selection

We evaluated the role of gene age in shaping the strength and pattern of selection on gene expression by employing a phylostratigraphy approach. In phylostratigraphy, proteomes of various species are aligned and underlying genes are partitioned into phylostrata (PS) dependent on the presence of homologs within particular clades. We downloaded PS values for each gene in the *C. elegans* genome from a recent study (Ma and Zheng 2023) and compared average selection differentials across each PS using Kruskal-Wallis tests (*kruskal_test())* followed by Dunn’s post-hoc tests (*dunnTest()*) in R.

### Population genomic information

We compared our inferred microevolutionary patterns of gene expression to rates of genome and protein coding sequence evolution to analyze how gene expression evolution relates to DNA and protein sequence evolution. Specifically, we retrieved *dN/dS* values from Ma *et al*. (2024), who calculated protein sequence divergence for *C. elegans* relative to 11 congeneric species. We also retrieved Tajima’s *D* values for *C. elegans* as determined by Lee *et al*. (2021) for 1-kb windows across the genome. We retrieved genomic locations for genes from the Ensembl WormBase ParaSite BioMart release 19 (https://parasite.wormbase.org/biomart/martview) *C. elegans* (PRJNA13758, WS290) dataset and assigned Tajima’s *D* values to genes based on the presence of its transcriptional start site within a given window.

We compared *dN/dS* and Tajima’s *D* values across genes of differing ages, tissue specificities, and network properties using the aforementioned statistical tests in R.

## Results

### Weak negative and stabilizing selection on the *C. elegans* transcriptome

To investigate the strength and pattern of natural selection on genome-wide gene expression in *C. elegans*, we leveraged transcriptome data from life stage-synchronized worms of a population of 207 genetically diverse strains curated by the *Caenorhabditis* Natural Diversity Resource (CaeNDR; Crombie *et al*. 2024). These strains represent species-wide genetic diversity and have been characterized by genome resequencing (Lee *et al*. 2021), genome-wide gene expression profiling (Zhang *et al*. 2022), and measurements of lifetime fecundity as a key fitness component (Zhang *et al*. 2021). Worms were reared in a common laboratory environment and collected for measuring transcript abundance during the onset of adulthood around the first embryo-laying event (Zhang *et al*. 2022), a critical time point for assessing reproductive fitness. Lifetime fecundity was calculated by summing progeny produced over four days by worms reared in the same laboratory environment (Zhang *et al*. 2021), providing an opportunity to infer direct links between gene expression patterns and fitness outcomes.

We applied genotypic selection analyses (Rausher 1992) to estimate selection differentials for 25,849 transcripts, representing approximately 16,094 genes. Two metrics were calculated: *S*, representing directional selection (whether transcript abundance covaries positively or negatively with fitness), and *C*, representing stabilizing or diversifying selection (selection for or against population-level variance in expression, respectively). After excluding 14 strains that produced no progeny under laboratory conditions, we calculated *S* and *C* for the remaining 193 strains.

Transcriptome-wide selection was weak overall, with *|S|_median_* = 0.018. Directional selection was biased towards negative selection (greater fitness with lower expression), with expression of 16,064 transcripts under negative selection (*S* < 0), and expression of 9,785 transcripts under positive selection, *i.e.*, greater fitness with higher expression (*S* > 0; *S_median_* = -0.2046 for transcripts under negative selection and *S_median_* = 0.01503 for transcripts under positive selection; Mann-Whitney *U*-test, two-sided, *p < 2.2×10^-16^*; Figure 1A). Stabilizing selection was the predominant form of selection with respect to population-level expression variance across the transcriptome, with 15,187 transcripts under stabilizing selection (*C* < 0) and 10,662 transcripts under diversifying selection, *i.e.*, selection for population-level variance (*C* > 0; *C_median_* = -0.022529 for transcripts under stabilizing selection and *C_median_* = 0.01535 for transcripts under diversifying selection; Mann-Whitney *U*-test test, two-sided, *p < 2.2×10^-16^*; Figure 1B). These findings corroborate prior evidence that weak stabilizing selection influences transcriptome evolution in *C. elegans* (Denver *et al*. 2005; Zhang *et al*. 2022; Bell *et al*. 2024). Full selection differentials for each transcript are reported in Supplementary Table 1.

**Figure 1:**
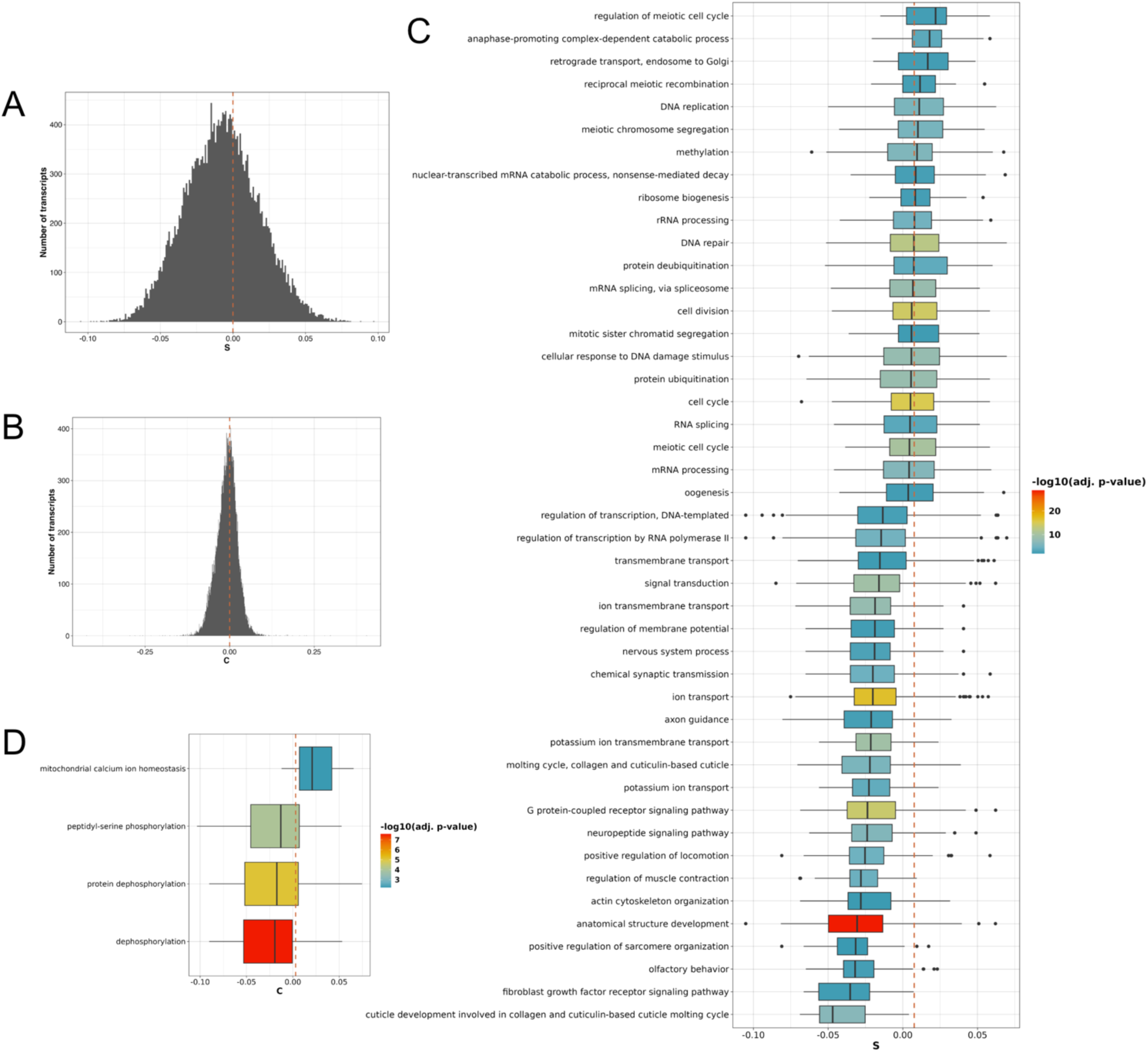
The strength and pattern of natural selection on transcript abundance in *Caenorhabditis elegans*. **A.** Distribution of linear selection differentials (*S*) on genome-wide gene expression. Vertical dotted orange line represents *S* = 0. **B.** Distribution of quadratic selection differentials (*C*) on genome-wide gene expression. Vertical dotted orange line represents *C* = 0. **C.** Gene Ontology (GO) biological processes with larger-than-average *S* values on the expression of underlying genes (Mann-Whitney *U*-test, two-sided, p < 1.15×10^-6^). Vertical dotted orange line represents the overall median selection differential *S*_median_ = 0.018. **D.** GO biological processes with larger-than-average *C* values on the expression of underlying genes (Mann-Whitney *U*-test, two-sided, p < 1.15×10^-6^). Vertical dotted orange line represents the overall median selection differential *C*_median_ = -0.00619.

Seven transcripts retained significant *S* values after Bonferroni correction (*p < 3.87 x 10^-7^*; Supplementary Figure S1). Among these, we observed significant positive selection on the expression of the UDP-glycosyltransferase *ugt-9*, which is known to be involved in detoxifying benzimidazole-like compounds (Sharma *et al*. 2024). Conversely, we observed negative selection on the expression of *B0348.2*, encoding a predicted zinc ion-binding protein; *Titin-1 (ttn-1)*, a key structural component of muscle filaments (Forbes *et al*. 2010); *feh-1*, an amyloid binding protein known to regulate pharyngeal pumping (Zambrano *et al*.); *clec-37*, a C-type lectin; *F25A2.1*, a gene with predicted contributions to lipid metabolism; and *nhr-114*, a nuclear hormone receptor responsive to nutritional cues (Gracida and Eckmann 2013; Giese *et al*. 2020; Qin *et al*. 2022). We will discuss potential relationships between variation in expression for several of these transcripts and fitness later. It was further notable that no transcripts retained significant *C* values after applying Bonferroni correction. Taken together, these results highlight that while selection acts weakly across most transcripts, few individual genes under stronger selection may have outsized contributions to fitness.

To examine selection on coordinated patterns of transcript abundance, we conducted PC analysis on genome-wide gene expression. The top 22 PCs, explaining at least 0.5% of transcriptome variation each, were subjected to multivariate regression to estimate direct linear (*β*) and quadratic (*Ɣ*) selection gradients (Supplementary Table 2). Among these PCs, eight exhibited significant directional selection (*β, p < 0.05*; Supplementary Figure 2A), and one showed significant diversifying selection (*Ɣ_PC1_ = 0.136, p = 0.0208*. Supplementary Figure 2B). However, after applying Bonferroni correction to our multivariate models (*p < 0.0022)*, only two PCs retained significant directional selection gradients (*β_PC4_ = 0.0858; p = 9.33×10^-8^; β_PC11_ = -0.04907; p = 0.00169*), while none retained significance for *Ɣ*. These results are concordant with the pervasive weak selection observed in our univariate analyses (Figure 1) and suggest that selection may target coordinated expression patterns rather than abundance of individual transcripts, consistent with theoretical models of gene network evolution.

To confirm that our selection estimates reflect biological processes rather than artifacts of population structure, we repeated the analysis after removing 21 strains identified as highly divergent relative to the other strains in the population based on genome-wide diversity metrics (Lee *et al*. 2021). The divergent strains exhibit lower relative fecundity in laboratory environments (Zhang *et al*. 2021), potentially confounding patterns of trait and fitness covariance. Exclusion of these strains did not significantly alter selection estimates, as indicated by high concordance between both models for *S* (Spearman *𝜌 = 0.9668, p < 2.2×10^-16^*) and *C* (Spearman *𝜌 = 0.9382, p < 2.2×10^-16^*) (Supplementary Figure S3, Supplementary Table 3).

Finally, we integrated our models with previously-identified QTL associated with total lifetime fecundity (Zhang *et al*. 2022). Of the 17 QTL, ten encoded at least one transcript with a significant linear selection differential (*S*), and two encoded a transcript significant for *C*, prior to Bonferroni correction (Supplementary Figure S4, Supplementary Table 4). Notably, *nhr-114* was among the transcripts overlapping with a fecundity-associated QTL and retained significance after Bonferroni correction (Supplementary Figure 1G). This integration reinforces the biological relevance of our findings and highlights potential regulatory mechanisms underlying fitness variation.

### Selection on gene expression is constrained by network architecture

To investigate how GRN architecture influences selection on gene expression, we integrated our selection models with information on eQTL mapped with the same transcriptome dataset (Zhang *et al*. 2022). Using the strength of selection (*|S|*) as a proxy for the direct (*i.e.*, microevolutionary) response of each transcript to selection (Lande and Arnold 1983; Hendry and Kinnison 1999), we compared average selection differentials across transcripts regulated by *cis-* and *trans-*eQTL.

We found that the strength of directional selection (*|S|*) increased with network connectivity. Expression of transcripts regulated by *cis*-eQTL experienced marginally stronger selection compared to expression of those without eQTL (Mann-Whitney *U*-test, two-sided, *p = 9.995×10^-5^*), and expression of transcripts regulated in *trans* experienced even stronger selection (for both *trans*-eQTL and *trans*-eQTL hotspots: Mann-Whitney *U-*test, two-sided, *p < 2.2×10^-16^*; Figure 2A). Stabilizing selection (*C < 0)* was slightly stronger for abundance of *cis*-regulated transcripts (Mann-Whitney *U*-test, two-sided, *p = 0.01119*), but selection on population-level variance in gene expression relaxed for transcripts regulated by *trans-*eQTL (Mann-Whitney *U*-test, two-sided, *p = 7.777×10^-4^*) and *trans*-eQTL hotspots (Mann-Whitney *U*-test, two-sided, *p = 0.03669*) (Figure 2B). These results suggest that expression of *cis*-regulated genes is subject to tighter regulatory constraints, while expression of *trans*-regulated genes experiences relaxed selective constraints.

**Figure 2:**
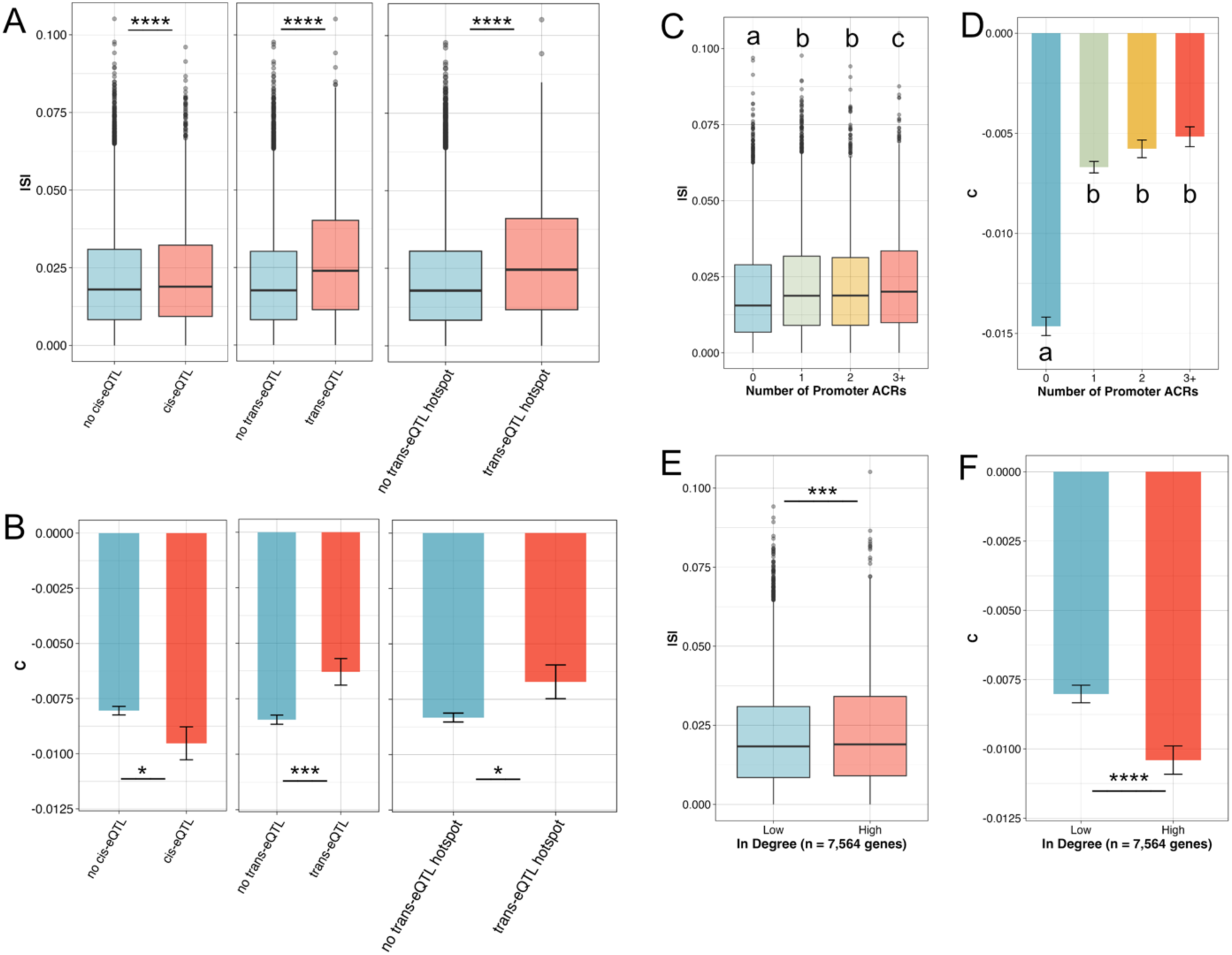
Network architecture constrains the strength of selection on gene expression. **A.** Relative to transcripts without eQTL, abundances of transcripts with eQTL, as mapped by Zhang *et al*. (2022), experience stronger directional selection (*cis*-eQTL: Mann-Whitney U-test, two-sided, p = 9.995×10^-5^; *trans*-eQTL: Mann-Whitney U-test, two-sided, p < 2.2×10^-16^; *trans*-eQTL hotspots: Mann-Whitney *U*-test, two-sided, p < 2.2×10^-16^). **B.** Relative to transcripts without eQTL, abundances of transcripts with *cis*-eQTL experience stronger stabilizing selection (Mann-Whitney *U*-test, two-sided, p = 0.01119), while stabilizing selection relaxes for transcripts regulated by *trans*-eQTL (Mann-Whitney *U*-test, two-sided, p = 0.000777) and *trans*-eQTL hotspots (Mann-Whitney *U*-test, two-sided, p = 0.03669). **C, D.** Expression of genes with more accessible chromatin regions (ACRs) experiences **C.** stronger directional selection (Kruskal-Wallis test with Dunn’s post-hoc test, p < 0.00266) and **D.** weaker stabilizing selection (Kruskal-Wallis test with Dunn’s post-hoc test, p < 9.734×10^-37^). **E, F.** Expression of genes regulated by more transcription factors experiences **E.)** stronger directional selection (Mann-Whitney *U*-test, two-sided, p = 0.0002268) and **F.** stronger stabilizing selection (Mann-Whitney *U*-test, two-sided, p = 2.644×10^-5^). For **B, D,** and **F,** columns represent mean *C* values and error bars represent standard error of the mean.

To link these findings to broader patterns of DNA and protein sequence evolution, we compared selection differentials to commonly-employed population genetic metrics. Namely, we retrieved genome-wide Tajima’s *D* values (Lee *et al*. 2021), which compare the number of segregating sites within a region of the genome to the average number of nucleotide differences across the species, providing insight into deviations from neutral evolution (when Tajima’s *D* is not 0). We also retrieved *dN/dS* values for each gene in the genome (Ma *et al*. 2024), which quantify relative rates of nonsynonymous to synonymous substitutions and serve as an indicator of the strength and direction of selection acting on protein-coding regions (neutral evolution when *dN/dS = 1*). *Cis-* regulated genes had average Tajima’s *D* values closer to 0 (Mann-Whitney *U*-test, two-sided, *p < 2.2×10^-16^*), suggesting relaxation of purifying selection on these genomic regions. In contrast, *trans-*regulated genes showed no significant deviation from genome-wide averages (*trans*-eQTL: Mann-Whitney *U*-test, two-sided, p = 0.5945; *trans*-eQTL hotspots: Mann-Whitney *U*-test, two-sided, p = 0.5634) (Figure 3A).

**Figure 3:**
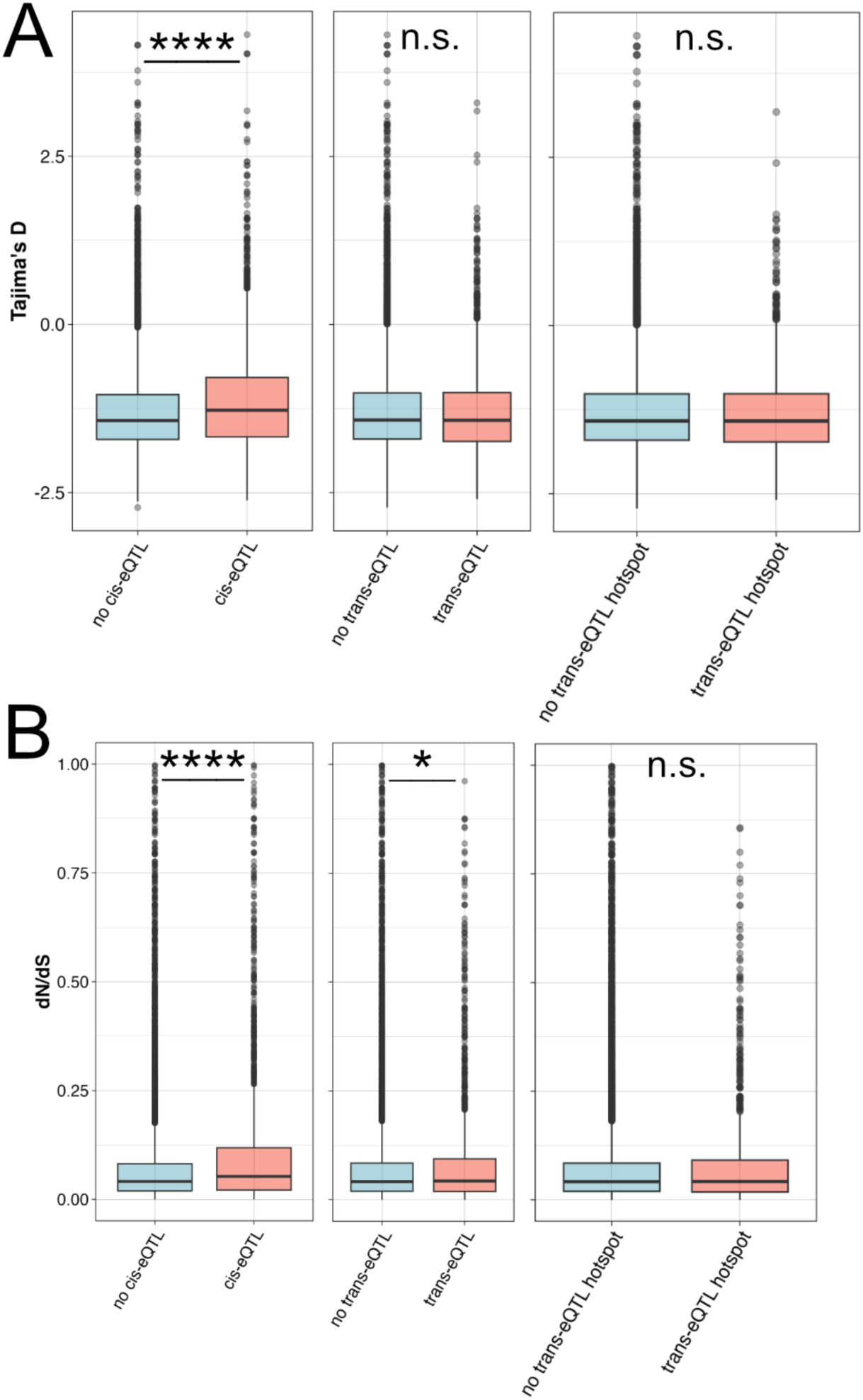
Rates of genome and protein sequence evolution are shaped by network architecture. **A.** Tajima’s *D* values for genes whose expression is regulated by eQTL. Relative to genes without eQTL, genes regulated by *cis*-eQTL experience slightly weaker purifying selection (Mann-Whitney *U*-test, two-sided, p < 2.2×10^-16^), but not those regulated by *trans*-eQTL (Mann-Whitney *U*-test, two-sided, p = 0.5945) or *trans*-eQTL hotspots (Mann-Whitney *U*-test, two-sided, p = 0.5634). **B.** *dN/dS* values for genes whose expression is regulated by eQTL. Relative to genes without eQTL, rates of protein sequence evolution are higher for genes regulated by *cis*-eQTL (Mann-Whitney *U*-test, two-sided, p < 2.2×10^-16^) and by *trans*-eQTL (Mann-Whitney *U*-test, two-sided, p = 0.0105), but not by *trans*-eQTL hotspots (Mann-Whitney *U*-test, two-sided, p = 0.199).

Genes regulated by *cis*-eQTL (Mann-Whitney *U*-test, two-sided, p < 2.2×10^-16^) and *trans*-eQTL (Mann-Whitney *U*-test, two-sided, p = 0.01054) also exhibited elevated *dN/dS* values, indicating higher rates of protein evolution. However, genes linked to *trans*-eQTL hotspots did not show this pattern (Mann-Whitney *U*-test, two-sided, p = 0.1999) (Figure 3B). These findings suggest that *cis*-regulated genes experience both rapid sequence evolution and strong stabilizing selection on expression, while *trans*-regulated genes are shaped by purifying selection, potentially to preserve network stability. The apparent relaxation of stabilizing selection on transcript levels for *trans*-regulated genes might facilitate network flexibility, enabling adaptive responses or accommodation for pleiotropy.

To further examine regulatory complexity, we incorporated tissue-specific chromatin accessibility data from L4-stage worms (Serizay *et al*. 2020). Expression of genes with more ACRs experienced stronger directional selection, regardless of whether chromatin was more accessible for promoter (Kruskal-Wallis test with Dunn’s post-hoc test, *p < 2.66×10^-^*^3^; Figure 2C) or enhancer elements (Kruskal-Wallis test with Dunn’s post-hoc test, *p < 2.732×10^-6^;* Supplementary Figure 5A). Stabilizing selection on gene expression was weaker for genes with more ACRs (for promoter ACRs: Kruskal-Wallis test with Dunn’s post-hoc test, *p < 9.734×10^-37^*; for enhancer ACRs: Kruskal-Wallis test with Dunn’s post-hoc test, *p < 0.0136*; Figure 2D; Supplementary Figure 5B). Despite these differences, Tajima’s *D* did not deviate significantly from genome-wide averages (Kruskal-Wallis test, *p = 0.3215*; Supplementary Figure 5C). Interestingly, genes with more ACRs exhibited lower *dN/dS* values (Kruskal Wallis test with Dunn’s post-hoc test, *p < 23.801×10^-26^*; Supplementary Figure 5D), suggesting that regulatory complexity enables fine-tuning of expression while maintaining sequence-level constraints.

We then mapped our selection differential models onto a GRN describing TF-target relationships derived from time-course transcriptomes of adult worms, which included timepoints surrounding the onset of adulthood (Suriyalaksh *et al*. 2022). Selection on the expression of TF-encoding genes did not differ by the out-degree of TFs (their number of target genes) for either *S* (Kruskal-Wallis test, *p = 0.3765*; Supplementary Figure 5E) or *C* (Kruskal-Wallis test, *p = 0.9406*; Supplementary Figure 5F). However, expression of genes with high in-degree (regulated by many TFs) experienced stronger directional selection (Mann-Whitney *U*-test, *p = 2.268×10^-4^*; Figure 2E) and stronger stabilizing selection (Mann-Whitney *U*-test, *p = 2.644×10^-5^*; Figure 2F).

Chi-square tests revealed significant overlap between high in-degree genes and those regulated by *trans*-eQTL (*χ*² = 62.474, p = 2.7×10⁻⁵) and *trans*-eQTL hotspots (*χ*² = 32.814, p = 1.0×10⁻¹⁴), but not *cis*-eQTL (*χ*² = 1.264, p = 0.2607). These position *cis*-eQTL at the periphery of GRNs and *trans*-eQTL closer to the core, consistent with previous findings (Mähler *et al*. 2017; Josephs *et al*. 2017). Despite their regulatory centrality, high in-degree genes showed no significant deviations in Tajima’s *D* (Mann-Whitney *U*-test, *p = 0.2324*; Supplementary Figure 5I) or *dN/dS (*Mann-Whitney *U-*test, p = 0.08082; Supplementary Figure 5J).

Our findings suggest that selection on gene expression is shaped by regulatory architecture and network position. Genes regulated by *cis*-eQTL evolve rapidly at the sequence level while maintaining constrained expression, whereas expression of *trans*-regulated genes experiences relaxed stabilizing selection. However, within a subset of *trans*-regulated genes, stabilizing selection on transcript level becomes stronger as in-degree increases, potentially reflecting the need to tightly constrain the expression of genes central to GRN stability. This relationship suggests that while *trans*-eQTL confer regulatory flexibility, high in-degree genes – often core components of GRNs – may impose constraints to buffer against the pleiotropic effects of expression variability. These results align with the omnigenic model, where diffuse regulation across GRNs drives complex trait variation, while central hubs ensure functional robustness.

### Tissue-specific transcriptomes evolve at different rates

To investigate how tissue specificity influences gene evolution, we calculated *τ* (tau) for each gene in the *C. elegans* genome. *τ* ranges from 0 (constitutive expression) to 1 (tissue-specific expression), providing a quantitative measure of expression breadth (Yanai *et al*. 2005). Using Fluorescence-Activated Nuclei Sorting (FANS)-RNA-sequencing data collected for four major tissues in L4-stage worms (Serizay *et al*. 2020), we found most genes are enriched in a limited number of tissues (Supplementary Figure 6A and Supplementary Table 5).

We observed that directional selection (|*S|*) was stronger for the expression of genes with higher tissue specificity (Kruskal-Wallis test with Dunn’s post-hoc test, *p < 6.581×10^-4^*; Figure 4A), and these genes also experienced stronger stabilizing selection on transcript levels (*C;* Kruskal-Wallis test with Dunn’s post-hoc test, *p < 9.299×10^-4^;* Figure 4B). While tissue-specific genes did not exhibit significant differences in Tajima’s *D* values (Kruskal-Wallis test, *p = 0.008209,* none significant after performing Dunn’s post-hoc tests; Supplementary Figure 6B), they showed higher rates of protein sequence evolution, as indicated by elevated *dN/dS* values (Kruskal-Wallis test with Dunn’s post-hoc test, *p < 2.310×10^-4^*; Supplementary Figure 6C). These findings align with prior observations that tissue-specific genes evolve faster at the protein level (Zhang and Li 2004), but might face tighter constraints on expression divergence (Liao and Zhang 2006).

**Figure 4:**
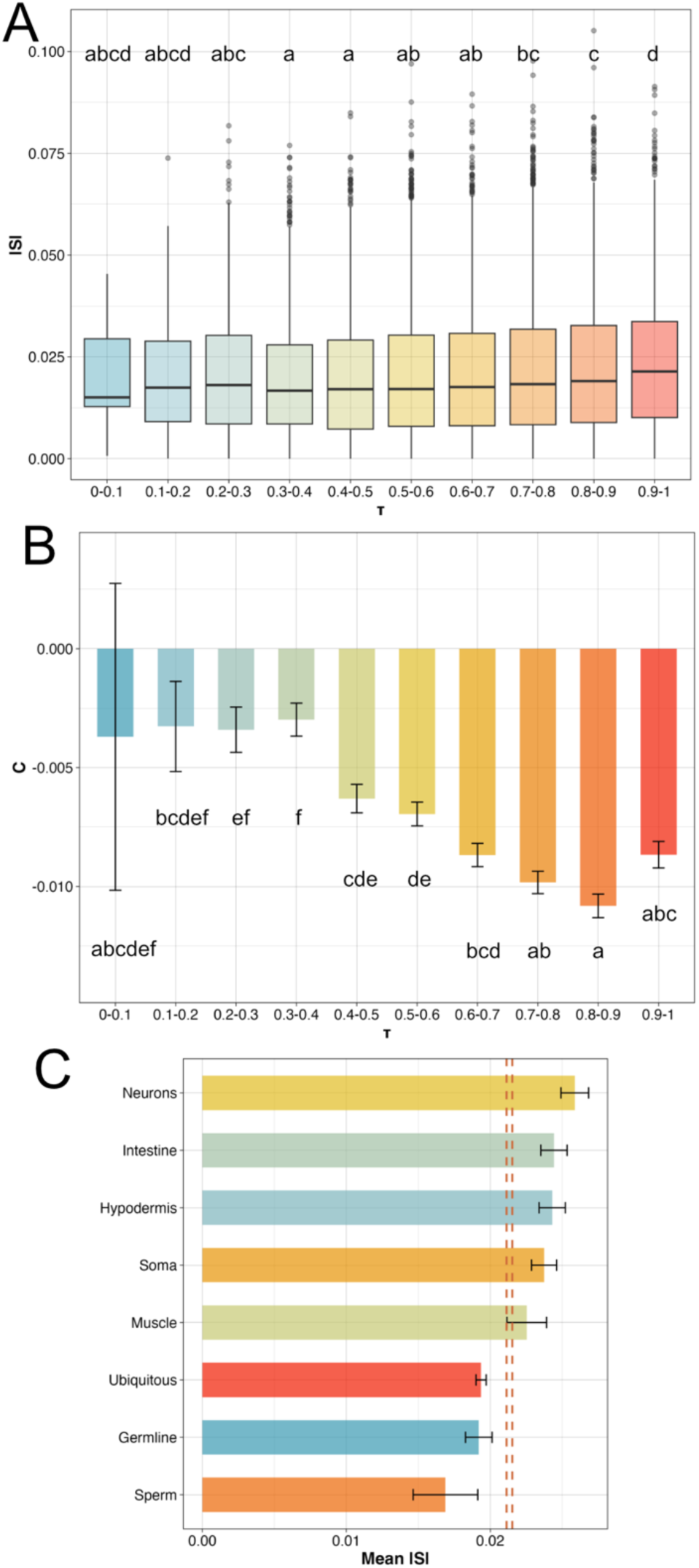
Tissue-specific transcriptomes experience stronger selection. **A, B.** Expression of genes with greater degrees of tissue-specificity (higher τ) experiences **A.** stronger directional selection (Kruskal-Wallis test with Dunn’s post-hoc test, p < 6.581×10^-4^) and **B.** stronger stabilizing selection (Kruskal-Wallis test with Dunn’s post-hoc test, p < 9.299×10^-4^). Columns represent mean *C* values and error bars represent standard error of the mean. **C.** Expression of germline- and sperm-specific genes experiences weaker selection, while expression of neuronal, intestinal, and hypodermal genes experiences stronger selection. Bars represent mean |*S*| values and error bars represent the 95% confidence intervals (CIs). Vertical dashed orange lines represent the 95% CIs for |*S*| across all genes in the genome.

Given the stronger directional selection observed for expression of tissue-specific genes, we assessed whether entire tissue-specific transcriptomes experience distinct selection pressures. We retrieved tissue-specific gene sets from Serizay *et al*. (2020) and calculated 95% confidence intervals for *|S|* for each tissue’s transcriptome (Figure 4C). Our analysis revealed that transcript abundances for neuronal, intestinal, hypodermal, and somatic tissue-specific genes experienced stronger-than-average directional selection, reflecting their critical roles in maintaining organismal fitness. Transcript abundances for muscle-specific genes showed values of *|S|* comparable to the genome-wide average, suggesting that selection on expression of these genes is not unusually strong or weak. In contrast, transcript abundances for germline-, sperm- and ubiquitously expressed genes experienced significantly weaker directional selection, suggesting a relative relaxation of selective constraints on these transcriptomes.

### Constraints on younger genes mirror constraints from network architecture and tissue-specificity

To investigate how gene age influences selection, we employed a phylostratigraphy approach, which assigns genes to approximate age classes based on the phylogenetic branch in which they first emerged (Domazet-Loso *et al*. 2007). Genes with lower phylostratum (PS) values represent older genes, whereas higher values indicate younger genes. We retrieved PS values from a recent study (Ma and Zheng 2023) and analyzed the strength of selection on expression of genes from different age classes.

Directional selection (|*S|*) was significantly stronger for expression of older genes, with a trend of increasing selection as PS values decreased (Kruskal-Wallis test with Dunn’s post-hoc test, *p < 6.812×10^-4^*; Figure 5A). The strongest selection was observed for expression of genes in PS2, corresponding to core ecdysozoan genes, while the weakest selection occurred for expression of PS10 genes, shared only between *C. elegans* and its close relative *C. briggsae*. PS2 genes were enriched for GO cellular compartments essential for neuromuscular function, such as the troponin complex, synaptic membrane, and neuron projections (Supplementary Figure 7A). These findings align with previous reports that younger genes are less likely to develop essentiality and contribute to fitness in *C. elegans* (Ma *et al*. 2024). They also support previous observations that laboratory environments impose strong selective pressure on neuromuscular systems in *C. elegans* (Sterken *et al*. 2015).

**Figure 5:**
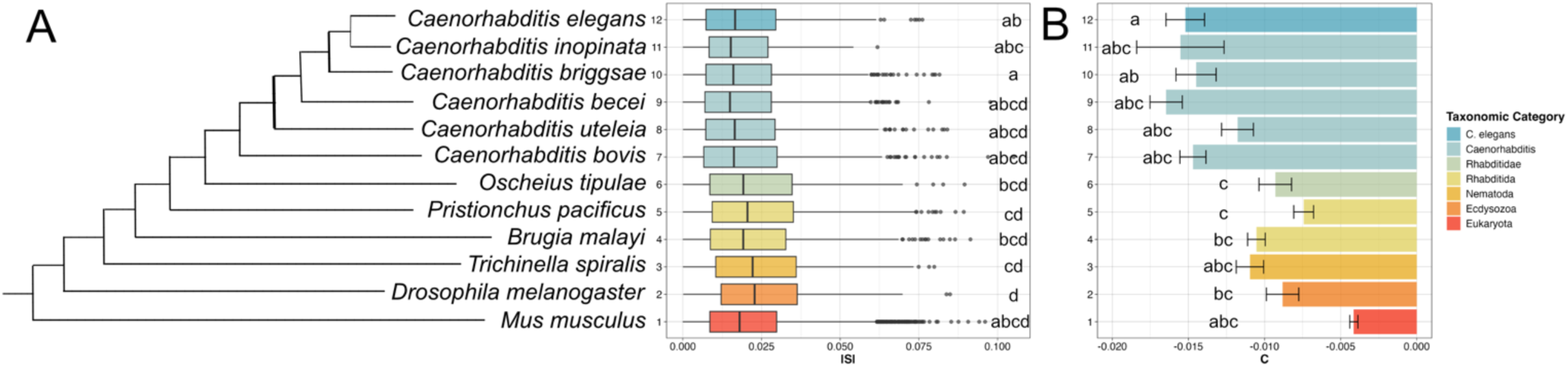
The landscape of selection is shaped by gene age. Genes were assigned relative ages, *i.e.*, they belonged to different phylostrata (PS1-12 along the y axes), based on their presence in orthogroups that were conserved in the phylogeny as in Ma and Zheng (2023). **A.** Expression of older genes experiences stronger directional selection (Kruskal-Wallis test with Dunn’s post-hoc test, p < 6.812×10^-4^), while **B.** expression of younger genes experiences stronger stabilizing selection (Kruskal-Wallis test with Dunn’s post-hoc test, p < 2.798×10^-4^).

Although younger genes experience relaxation of purifying selection at the coding sequence level (Ma *et al*. 2024), we found that stabilizing selection (*C*) was strongest for expression of younger genes (Kruskal-Wallis test with Dunn’s post-hoc test, *p < 2.798×10^-4^*; Figure 5B). This suggests that constraints on expression evolution might decrease as genes age, reflecting a shift in selective pressures over time.

We hypothesized that the stronger stabilizing selection on expression of younger genes might reflect their tendency to adopt tissue-specific expression profiles (Fukushima and Pollock 2020; Ma *et al*. 2024) and integrate into the periphery of GRNs (Capra *et al*. 2010; Zhang *et al*. 2015; Wei *et al*. 2016; Defoort *et al*. 2018). Consistent with this hypothesis, younger genes exhibited higher tissue-specificity (Kruskal-Wallis test with Dunn’s post-hoc test, *p < 3.022×10^-4^*; Supplementary Figure 7B), and were associated with fewer ACRs (Kruskal-Wallis test with Dunn’s post-hoc test, *p < 4.685×10^-4^*; Supplementary Figures 7C-D). These findings suggest that younger genes may face initial constraints that limit their expression flexibility, facilitating their integration into specific tissues and peripheral network roles.

### Functional enrichment of univariate and multivariate selection models

To uncover the biological processes driving these patterns of selection on transcript abundance, we examined the distribution of selection differentials across GO categories. Using Mann-Whitney *U*-tests, we identified GO terms with distinct distributions of *S* and *C* values, highlighting functional categories where transcript abundance covaries uniquely with fitness. While *|S|* provides a valuable measure of the overall strength of selection, it does not capture the direction of selective pressures - whether gene expression covaries positively or negatively with fitness. By unfolding *|S|* into *S*, we aimed to better resolve how transcript abundance aligns with fitness across functional categories.

Following Bonferroni correction (*p < 1.15×10^-6^*), we identified 45 GO biological processes with stronger-than-average positive or negative directional selection (*S*) on expression of underlying genes (Supplementary Table 6). Expression of suites of genes associated with reproduction-associated GO terms, such as regulation of the meiotic cell cycle, reciprocal meiotic recombination and meiotic chromosome segregation tended to experience stronger positive selection, while expression of suites of genes associated with behavior- and neuromusculature-associated GO terms, including chemical synaptic transmission, axon guidance, neuropeptide signaling and regulation of muscle contraction, tended to experience stronger negative selection (Figure 1C). These patterns align with prior findings suggesting that key laboratory-derived alleles in the *C. elegans* reference strain N2 enhance fitness relative to wild strains in laboratory environments by accelerating egg production rates (Duveau and Félix 2012) and by repressing neurological processes linked to social, sensory, and feeding behaviors (de Bono and Bargmann 1998; Rogers *et al*. 2003; Gray *et al*. 2004; Chang *et al*. 2006; McGrath *et al*. 2009; Macosko *et al*. 2009; Bendesky *et al*. 2011; Andersen *et al*. 2014). They further suggest that the “laboratory domestication” of *C. elegans* might involve broad transcriptional reprogramming (Rockman *et al*. 2010; Andersen *et al*. 2014; Sterken *et al*. 2015), mirroring patterns observed in domesticated crops such as maize (*Zea mays;* Swanson-Wagner *et al*. 2012) and tomato (*Solanum lycopersicum*; Koenig *et al*. 2013).

When we repeated these analyses for *C*, we found four GO biological processes whose suites of associated genes experienced stronger-than-average selection on population-level variance in expression (Supplementary Table 7). Among these, suites of genes involved in modulating kinase signaling networks, such as peptidyl-serine phosphorylation and protein dephosphorylation, exhibited stronger stabilizing selection on their expression, suggesting tight regulation to maintain fitness. In contrast, suites of genes associated with mitochondrial calcium ion homeostasis experienced stronger diversifying selection on their expression, reflecting population-level variability that may support adaptive responses to environmental variation (Figure 1D).

We further tested for enrichment of GO biological processes among genes loading strongly into PC3 and PC4 from our PC analysis because these PCs explain a relatively large proportion of transcriptome variation and experience direct positive selection (*β > 0*; Figure S2A). Among the top 100 transcripts loading into either PC3 or PC4, we observed enrichment of biological processes related to RNA processing, such as ribosomal RNA (rRNA) processing, non-coding RNA (ncRNA) metabolic processes, and mitochondrial translation (Supplementary Figure S2C-D). Notably, this enrichment aligns with the results of our univariate analyses, which similarly revealed positive selection on the expression of various components of RNA processing pathways (*S > 0*; Figure 1C). These concordant results suggest that pathways regulating RNA metabolism may play a central role in determining organismal fitness.

### Selection on gene expression highlights trade-offs between reproductive strategies

After unfolding *|S|*, we revisited our analysis of tissue-specific gene sets and found that transcript abundance for germline-specific and ubiquitously expressed genes experienced stronger-than-average positive selection, while transcript abundance for genes expressed in all other tissues experienced relatively stronger negative selection (Figure 6A). This pattern closely mirrors the results of our GO analysis, reinforcing the notion that investment in reproduction-associated gene expression positively influences fitness, whereas investment in gene expression related to organismal physiology negatively affects fitness, at least during the adolescent transition (Figure 1C).

**Figure 6:**
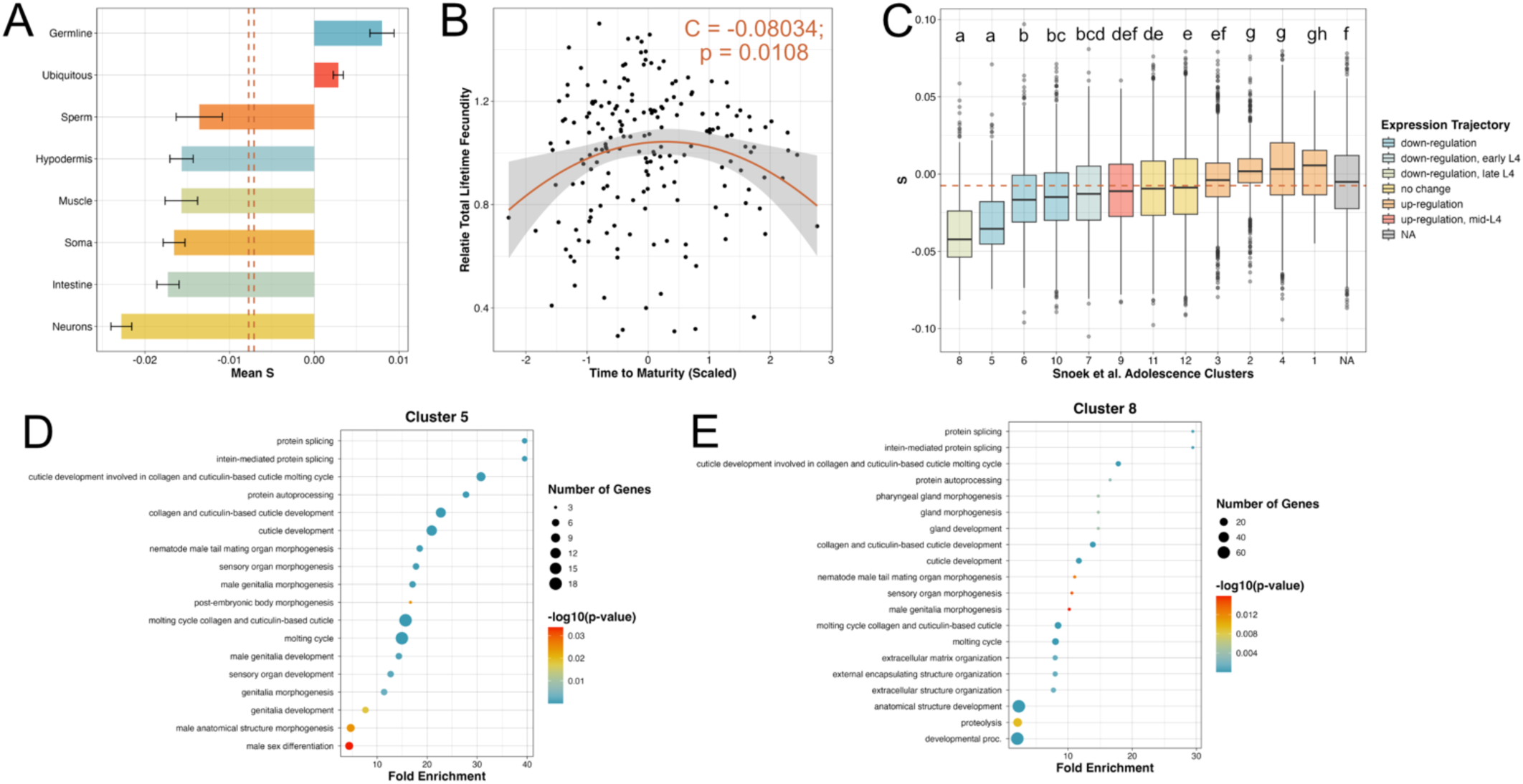
Selection mediates trade-offs between nematode reproductive strategies. **A.** Expression of germline-specific and ubiquitous genes experiences positive selection, while expression of all other tissue-specific genes experiences negative selection. Bars represent mean *S* values with 95% confidence intervals (CIs). Vertical dashed orange lines represent 95% CIs for *S* across all genes in the genome. **B.** Covariation between time to maturity and fecundity reveals weak stabilizing selection on developmental timing in *C. elegans* (*C* = -0.0803 ± 0.0312, p = 0.0108). **C.** Selection on expression of coregulated clusters of transcripts with dynamic expression patterns during the adolescence transition as determined by Snoek *et al*. (2014). While most clusters have mean *S* values near the genome-wide median selection differential (horizontal dashed orange line; *S* = -0.018), expression of genes in clusters 5 and 8 experiences stronger negative selection (Kruskal-Wallis test with Dunn’s post-hoc test, p < 2.746×10^-4^). **D, E.** Results of Gene Ontology biological process enrichment for genes in **D.** cluster 5 and **E.** cluster 8 suggest that selection reduces the expression of genes that promote the development of male-specific traits.

Most intriguing was the finding that, despite positive selection on the expression of germline-specific genes, expression of sperm-specific genes experienced negative selection. We hypothesize that this pattern reflects a reproductive conflict in *C. elegans*, where spermatogenesis and oogenesis are temporally separated, with an irreversible shift to oogenesis occuring at the onset of fertilization (Ward and Carrel 1979). While models suggest that sperm production limits lifetime fecundity in *C. elegans* under optimal conditions (Cutter 2004), experimental evidence indicates that stressful environments may mitigate these constraints (Murray and Cutter 2011). To explore further, we reapplied genotypic selection analysis to test for covariance between fitness and time to maturity – a key determinant of sperm production (Cutter 2004; Murray and Cutter 2011) – predicted by strain-specific transcriptome data (Zhang *et al*. 2022). Although we detected no significant linear relationship between time to maturity and fecundity (*S = 0.002955 ± 0.018596, p = 0.874*; Supplementary Figure S8), weak but significant stabilizing selection was observed (*C = -0.08034 ± 0.0312, p = 0.0108*; Figure 6B). These findings suggest that the global *C. elegans* population is near a fitness optimum for maturation timing, where trade-offs between sperm and egg production are modulated by generation time. Increased investment in sperm production likely enhances fitness only up to a critical threshold, beyond which extended generation times incur fitness costs (Cutter 2004; Murray and Cutter 2011).

To identify the genes mediating these trade-offs in maturation timing, we examined genes with dynamic expression patterns during adolescence (Snoek *et al*. 2014). Two clusters of genes whose expression experienced the strongest negative selection on average were identified (Kruskal-Wallis test with Dunn’s post-hoc test, *p < 2.746×10^-4^*; Figure 6C). These clusters were enriched for GO terms associated with molting (*e.g*., cuticle development, molting cycle, etc.) and the development of male-specific sexual characteristics (*e.g*., male genitalia development, nematode male tail mating organ morphogenesis; Figures 6D-E). This pattern reinforces the notion that investment in male-specific traits incurs fitness costs, leading to selection against them in laboratory environments.

### Selection might act on the activity of transcription factors

We hypothesized that selection might act on the activity of TFs instead of or in addition to acting on the expression of the genes encoding them. To address this hypothesis, we examined whether groups of TF targets experienced stronger-than-average selection on their transcript levels. We retrieved both experimentally validated and computationally predicted TF-target relationships curated by Suriyalaksh *et al*. (2022). After Bonferroni correction (*p < 2.517×10^-5^*) we identified 19 TFs as candidates whose activity might be targeted by selection (Figure 6A).

Of these, 10 had annotated mutant or RNA interference phenotypes in WormBase linked to reproductive output (*e.g*., sterile, reduced brood size, egg-laying defective, fewer germ cells, etc.), underscoring their potential contributions to organismal fitness. For example, expression of the *EGg-Laying defective 38 (egl-38)* regulon experienced stronger-than-average positive selection (Figure 7A). *Egl-38* encodes a paired box (Pax) TF that mediates patterning of the uterine lining and vulva and regulates the expression of key neuropeptides involved in the timing of egg laying (Chamberlin *et al*. 1997; Webb Chasser *et al*. 2019). Mutants of *egl-38* are unable to lay eggs and accumulate germline cell corpses, reflecting its essential role in reproduction (Trent *et al*. 1983; Park *et al*. 2006; Rajakumar and Chamberlin 2007).

**Figure 7:**
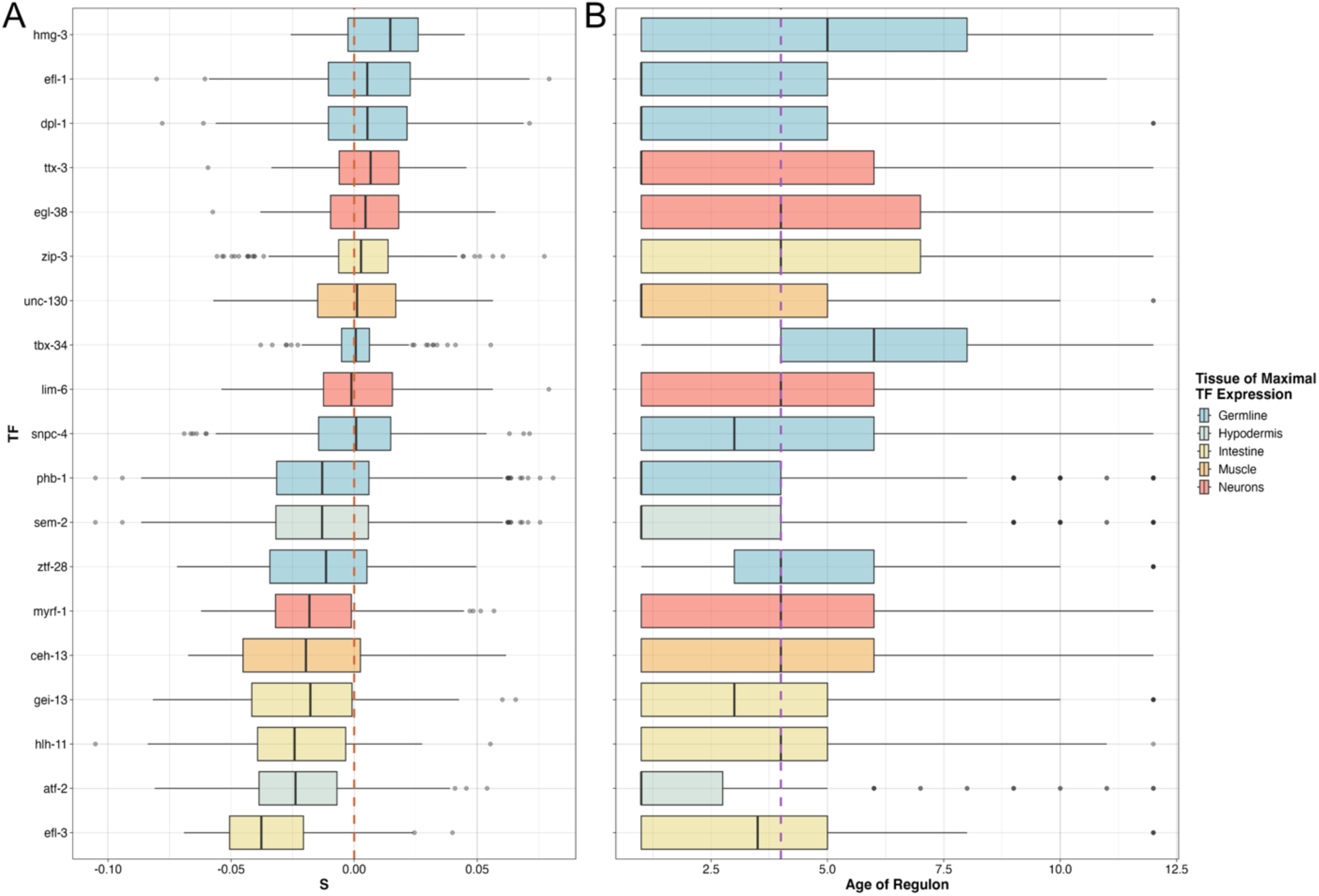
Selection acts on the activity of a subset of transcription factors. **A.** Nineteen transcription factors (TFs) have targets whose average selection differentials (*S*) for their expression diverge from the null distribution (Mann-Whitney *U*-test, two-sided, p < 2.517×10^-5^). Vertical dashed orange line at *S* = 0. **B.** Average age of each TF’s regulon as determined by the phylostrata values determined by Ma and Zheng (2023) and depicted in Figure 4. Vertical dashed purple line represents the median age of all genes in the genome (PS = 4). Most TFs with stronger-than-average selection on their activity regulate the expression of relatively old genes. Boxes are colored by the tissue in which each TF exhibits highest expression as determined by Serizay *et al*. (2020).

Given that expression levels of older (Figure 5) and tissue-specific (Figure 4) genes experienced relatively strong positive selection, we reasoned that these TFs might exhibit tissue-specific enrichment and regulate suites of older genes. Consistent with the observation that muscle-enriched genes experienced average levels of selection on their expression, activity of only two of the 19 TFs was enriched in the musculature (Figure 7A). By contrast, activity of the highest proportion (8/19) of the TFs was enriched in the germline, followed by activity of those enriched in neurons (4/19) and the intestine (4/19), aligning with these tissues’ transcriptomes exhibiting the strongest selection differentials when considering both *S* and *|S|* (Figures 4C, 6A).

To further explore the evolutionary dynamics of these TFs, we approximated the age of each TF’s regulon by calculating the average PS value across its targets. While the genome-wide average age class for a gene was PS4, half of the regulons (10/19) had average PS values less than four, suggesting that these TFs might preferentially regulate older genes (Figure 7B). This finding supports the hypothesis that selection acts on TF activity to fine-tune the expression of evolutionarily conserved genes critical for fitness, particularly in germline and neuronal tissues.

## Discussion

Understanding how patterns of gene expression evolve and what features of GRNs contribute to their evolutionary dynamics is a central question in evolutionary biology (Carroll 2008; Romero *et al*. 2012). However, most changes in gene expression have no significant impact on fitness and emerge as a consequence of genetic drift (Lynch 2007). Methodologies that can discriminate between adaptive evolution and genetic drift are essential for uncovering the selective forces shaping GRNs and provide critical insights into the genetic basis of phenotypic diversity (Price *et al*. 2022). By quantifying associations between transcript abundance and fecundity, we were able to identify gene expression patterns that are likely to be adaptive in *C. elegans* (Figure 1).

### Evidence for adaptive gene expression patterns

After implementing multiple testing corrections, seven transcripts retained significant univariate selection differentials (*S;* Supplementary Figure S1). Among these was one derived from the gene encoding mammalian *FE65 Homolog 1* (*feh-1)*, whose abundance covaried negatively with lifetime fecundity (Supplementary Figure S1D). *Feh-1* encodes the amyloid precursor beta binding protein, which is implicated in Alzheimer’s disease (Zambrano *et al*. 2002). In *C. elegans,* it is expressed in the pharyngeal neuromusculature, where it regulates pumping rates in a dose-dependent manner (Zambrano *et al*. 2002). Since pharyngeal pumping is a key feature of feeding behaviors (Doncaster 1962; Bhatla *et al*. 2015), it is intriguing to hypothesize that reduced *feh-1* expression in the most fit strains may enhance feeding efficiency, thereby supporting higher reproductive success.

Another notable example was a transcript derived from the nuclear hormone receptor-encoding gene *nhr-114*, whose abundance exhibited the strongest and most significant negative covariation with fecundity in the genome (Supplementary Figure S1G). *Nhr-114* is induced during vitamin B12 deficiency and participates in feedback loops that protect germline stem cells from oxidative damage (Gracida and Eckmann 2013; Giese *et al*. 2020; Qin *et al*. 2022). Variation in *nhr-114* expression might reflect population-level differences in nutritional status or in altered feeding behaviors that arise in laboratory environments (Sterken *et al*. 2015). These examples underscore the potential for nutrient-sensing and metabolic regulation pathways to make adaptive contributions to fitness, particularly in environments with novel selective pressures.

### Weak selection across the transcriptome

Apart from the seven transcripts identified through our selection analyses, the strength of selection was generally weak, as reflected in both the linear (*S* and *β*, Figure 1A and Supplementary Figure 2A) and quadratic (*C* and *Ɣ*, Figure 1B and Supplementary Figure 2B) selection differentials and gradients. This finding aligns with theoretical models of selection on complex traits, which predict that most phenotypic variation is subject to weak selective pressures due to polygenic architectures and pleiotropy (Kingsolver *et al*. 2001; Hoekstra *et al*. 2001). It is also consistent with observations in other systems, including ones of weak directional selection on transcript abundance in rice, salmonid fish, and the red flour beetle *Tribolium castaneum* (Groen et al. 2020; Koch and Guillaume 2020; Ahmad et al. 2021), and of trends towards weak stabilizing selection (*C < 0*) in rice (Groen *et al*. 2020). Together, these suggest that weak selection on gene expression is a common feature across species on microevolutionary timescales and may reflect the inherent, scale-free properties of GRNs that stabilize transcriptomes through regulatory complexity and buffering mechanisms (Whitacre and Bender 2010; Macneil and Walhout 2011; Whitacre 2012).

Despite the generally weak selection on shorter timescales, population averages in transcript abundance remain capable of shifting over time. This is supported by the observation that approximately one-fifth of the transcripts measured in the *C. elegans* population that constitutes the CaeNDR panel exhibit high SNP-based heritability (h^2^; Zhang *et al*. 2022) Subsets of these transcripts further experience relatively strong directional selection, which is related to their regulation by various eQTL (Figure 2). Their high heritability indicates that genetic variation in the abundance of these transcripts has potential to respond to selection over multiple generations, highlighting the latent capacity of gene networks to evolve, even when selective pressures are diffuse.

### Tissue-specific patterns of selection

Tissue-specific genes are associated with elevated rates of protein sequence evolution (Zhang and Li 2004) and tissue-specific transcriptomes diverge at different rates over long evolutionary timescales (Brawand *et al*. 2011; Fukushima and Pollock 2020; El Taher *et al*. 2021; Mantica *et al*. 2024). However, whether they evolve at different rates on microevolutionary timescales has remained unclear. Using *τ* as a metric of tissue-specific gene expression, we found that tissue-specific genes experience stronger-than-average directional selection on their expression (|*S*|; Figure 4A), suggesting that these genes have the potential to evolve faster at the expression level. This observation agrees with positive correlations between *|S|* and *τ* in rice (Groen *et al*. 2020). Additionally, tissue-specific genes exhibited stronger stabilizing selection on their expression (*C*; Figure 4B). This suggests the existence of constraints related to their regulatory sequences, which is consistent with findings in mammals (Liao and Zhang 2006).

Among tissue types, the strength of selection (*|S|*) was unexpectedly high for the expression of neuronal genes, possibly reflecting the unique selective pressures laboratory environments impose on the nervous system (Figure 4C; Sterken *et al*. 2015). In contrast, the relatively strong selection on the expression of intestinal genes draws parallels to findings in vertebrates, where analogous organs involved in detoxification, such as the kidneys and liver, evolve rapidly (Brawand *et al*. 2011; El Taher *et al*. 2021; Mantica *et al*. 2024). Expression of germline genes experienced the weakest absolute selection (*|S|*; Figure 4C), but notably, they were the only tissue-specific genes whose expression experienced positive selection (*S > 0*; Figure 6A), aligning with the rapid evolution of reproductive transcriptomes reported across diverse taxa (True and Haag 2001; Brawand *et al*. 2011; Fukushima and Pollock 2020; El Taher *et al*. 2021; Mantica *et al*. 2024). Together, these findings suggest that the principles governing tissue-specific GRN evolution over longer timescales may shape microevolutionary patterns of gene expression as well.

We found that expression of suites of genes involved in meiosis and oogenesis was under positive selection (Figure 1C), suggesting that strains with higher fitness might exhibit faster rates of egg production. This aligns with genotypic selection analyses on gene expression in *Tribolium castaneum,* where relatively strong positive selection was observed on *Vitellogenin* genes that produce precursors to egg yolk proteins (Koch and Guillaume 2020). These observations suggest that egg production rates may be a key determinant of fecundity across ecdysozoans. Supporting this, we identified TFs whose activity appears to be under selection, with several of them regulating genes directly involved in reproduction. For example, the regulon of *egl-38*, a Pax TF critical for uterine and vulval development and the timing of egg-laying (Chamberlin *et al*. 1997; Webb Chasser *et al*. 2019), was under positive selection for its expression (Figure 7A). This focus of selection on egg production and laying appears to contradict theoretical models predicting that rates of sperm production, rather than egg production, should limit fecundity in *C. elegans* (Cutter 2004). However, our results show weak stabilizing selection on time to maturity (Figure 6B) and negative selection on the expression of genes regulating the development of male sexual characteristics (Figures 6C-D). This suggests that in relatively novel laboratory environments, strains may optimize fitness by balancing investment in both egg- and sperm production. This interpretation aligns with previous findings that suboptimal or stressful environments may reverse sperm limitation (Murray and Cutter 2011).

### The nervous system as a focal point of evolution

In addition to the germline, we observed consistent and distinct patterns of selection on expression of genes active in the nervous system. Specifically, selection appeared to favor lower expression of genes involved in neurophysiology (Figure 1C), suggesting that reduced expression of these genes may enhance fitness in laboratory environments. Neuron-enriched genes exhibited the strongest directional selection on expression across tissue-specific transcriptomes (|*S*|; Figure 4C), which we determined to be predominantly due to negative selection on expression levels (*S < 0*; Figure 6A). Additionally, the stronger-than-average directional selection on the relatively old genes in PS2 was largely driven by genes whose expression was localized to neurons and synapses (Supplementary Figure S7A). These patterns suggest that laboratory environments might impose strong evolutionary constraints on neurophysiological pathways. Supporting this interpretation, several key laboratory-associated alleles in the reference strain N2 are known to reshape the nervous system. As an example, N2 possesses a gain-of-function mutation in the neuropeptide receptor-encoding gene *npr-1*, which expands its ligand binding repertoire relative to wild strains, effectively repressing neural processes that signal through the RMG interneuron (Rogers *et al*. 2003; Macosko *et al*. 2009). This mutation has been linked to successful adaptation to laboratory environments by reducing the aerotaxis response, which causes wild strains to clump on agar plates, reducing fitness by creating resource competition and starvation (de Bono and Bargmann 1998; Rogers *et al*. 2003; Gray *et al*. 2004; McGrath *et al*. 2009; Macosko *et al*. 2009). These observations illustrate how selective pressures unique to controlled environments can drive significant changes in neural gene regulation and behavior by optimizing neural functions to match environmental demands. Such adaptations underscore the broader role of the nervous system as a focal point of evolutionary change in response to novel ecological pressures (Cande *et al*. 2013; Konstantinides and Desplan 2024).

### Network architecture and regulatory variation

Network position is believed to constrain rates of evolution because core genes share pleiotropic relationships with many others, enhancing the potential fitness cost of mutations that occur within them (He and Zhang 2006). Consistent with this principle, we observed stronger directional selection (*S*) on the expression of core, high in-degree genes (Figure 2E). Additionally, expression of these genes was subject to stronger stabilizing selection (*C < 0*; Figure 2F), suggesting that selection acts to preserve network integrity by purging expression variance of their cores. Interestingly, we observed a reduction in the strength of stabilizing selection for the expression of genes with greater chromatin accessibility (Figure 2D, Supplementary Figure 5B). At first glance, this contrast may seem paradoxical; however, even ubiquitously expressed genes often possess distinct subsets of ACRs in different tissues (Serizay *et al*. 2020). This suggests that while high chromatin accessibility might overlap with high in-degree genes, it may also indicate compensatory or redundant regulatory mechanisms that buffer expression of these genes from the full impact of stabilizing selection.

The relationship between network position, regulatory variation, and selection is further supported by comparative studies in *Capsella grandiflora* and *Populus tremula*. In these plant species, genes regulated by *cis*-eQTL tend to exhibit lower connectivity and are restricted to the periphery of expression networks (Mähler *et al*. 2017; Josephs *et al*. 2017). Conversely, *trans*-regulated genes are more likely to occupy network cores in *P. tremula* (Mähler *et al*. 2017). These patterns align with our observation that *trans*-regulated transcripts overlap with high in-degree genes, while *cis*-regulated transcripts do not. The elevated Tajima’s *D* and *dN/dS* levels for genes regulated by *cis-*eQTL in *C. elegans* (Figure 3) are also consistent with findings in *C. grandiflora* and *P. tremula*, suggesting that local regulatory variation weakens the effects of purifying selection, allowing for more rapid evolution at the network periphery (Mähler *et al*. 2017; Josephs *et al*. 2017).

However, when comparing selection on expression between genes with and without different types of eQTL in the context of GRNs, we find that the abundance of *cis*-regulated transcripts in *C. elegans* experiences stronger stabilizing selection, while levels of *trans*-regulated transcripts exhibit relaxed stabilizing selection (Figure 2B). This may indicate that core genes, while generally under strong stabilizing selection, harbor *trans*-acting sequence variants that reduce expression variance constraints to maintain diversity in a subset of their regulatory elements. Conversely, *cis*-regulatory variation may facilitate the integration of genes into the periphery of expression networks, where they contribute to evolutionary innovation. Notably, recently duplicated and species-specific genes are enriched for eQTL in *P. tremula* (Mähler et al. 2017), drawing parallels to our observation of increased stabilizing selection on expression of younger genes (Figure 5B). Together, this suggests that while network architecture limits expression evolution by imposing selective constraints on core components, key genetic variants, such as certain *cis-* and *trans-* regulatory SNPs, may shift these constraints. This interplay between network structure and regulatory variation underscores how genetic mechanisms might modulate the balance between robustness and adaptability in GRNs.

### Gene age and GRN integration

Directional selection (*|S|*) was unexpectedly strong for expression of older genes (Figure 5A), largely driven by positive selection on expression of neuromusculature-related genes (Supplementary Figure S7A), suggesting that the selective constraints imposed on nervous systems by laboratory environments might override correlations with gene age. In contrast, stabilizing selection (*C < 0*) dominated among expression of genes in younger phylostrata, with its strength diminishing for genes that predate the divergence of the genus *Caenorhabditis* (Figure 5B). We hypothesize that elevated stabilizing selection on expression of genes in younger phylostrata reflects their tissue specificity (Supplementary Figure S7B) and ongoing integration into GRNs (Supplementary Figures S7C-D). This aligns with studies suggesting that gene expression divergence between species often follows an Ornstein-Uhlenbeck (OU) process, where between-species variance plateaus rapidly due to stabilizing selection, underscoring its central role in transcriptome evolution (Bedford and Hartl 2009; Nourmohammad *et al*. 2017; Chen *et al*. 2019). Our findings support these models, since purging of population-level expression variance as expression nears its fitness optimum may explain the reduced stabilizing selection on expression observed for older genes, potentially reflecting their more established roles within GRNs.

### TF activity and coordinated selection

We identified 19 TFs whose regulons are under relatively strong positive or negative selection on their expression (Figure 7A), many of which have been empirically linked to fecundity. Consistent with stronger directional selection observed for expression of tissue-specific (Figure 4) and older (Figure 5) genes, we found that many of these TFs are enriched in tissues experiencing relatively strong selection on gene expression, such as the germline, neurons, and intestine (Figure 7B). These TFs also regulate suites of older genes, potentially explaining the observed correlations between tissue specificity, gene age, and strength of selection on transcript abundance (Figure 7B). Together, these findings suggest that natural selection fine-tunes gene expression through the coordinated activity of key TFs, shaping the regulatory landscape of tissues critical for fitness.

Interestingly, even TFs without defined links to fecundity emerged as promising candidates for functional exploration. For example, we observed positive selection on the regulon of the bZIP TF *zip-3*. Along with its paralog *atfs-*1, this TF is a key regulator of the mitochondrial unfolded protein response (UPR; Fiorese *et al*. 2016; Deng *et al*. 2019). Unlike *atfs-1*, *zip-3* functions as a negative regulator, preventing hyperactivation of the mitochondrial UPR (Deng *et al*. 2019). Given that mitochondrial dysfunction is a conserved hallmark of aging (Brys *et al*. 2010; López-Otín *et al*. 2013), and is tightly linked to aging-related transcriptional changes in *C. elegans* (Roux et al. 2023), *zip-3* may play a role in mediating population-level differences in fecundity by balancing stress responses tied to mitochondrial energy status. Future studies investigating *zip-3* activity across strains could reveal how mitochondrial function intersects with reproduction and stress resilience.

## Conclusion

Our findings highlight how the strength and patterns of selection on gene expression in *C. elegans* are shaped by GRN architecture, tissue specificity, and regulatory variation. While network constraints impact stabilizing selection on the expression of core genes, regulatory variation and chromatin dynamics provide opportunities for adaptive shifts, balancing robustness with flexibility. These results suggest that the principles governing GRN evolution over macroevolutionary timescales also operate at microevolutionary timescales, shaping phenotypic diversity through the interplay of genetic and regulatory mechanisms.

## Data Availability

Normalized genome-wide gene expression and total lifetime fecundity values across 207 *C. elegans* strains were retrieved from Supplementary Data File 1 from Zhang *et al*. (2022). Expression QTL data were retrieved from Supplementary Data File 2 from Zhang *et al*. (2022). Chromatin accessibility information, tissue-specific gene expression values, and tissue-specific gene sets were retrieved from Supplementary Table 2 from Serizay *et al*. (2020). For our GRN, we used the “max AUFE” model from Table S6A from Suriyalaksh *et al*. (2022) after filtering out non-TF regulators, including the insulin receptor *daf-2*, and chromatin cofactors. Phylostratigraphy values were retrieved from Table S1 from Ma and Zheng (2023). *dN/dS* values for *C. elegans* were retrieved from Table S4 from Ma *et al*. (2024). Tajima’s *D* values across the *C. elegans* genome as calculated by Lee *et al*. (2021) were downloaded from GitHub at https://github.com/AndersenLab/Ce-328pop-div.

## Acknowledgments

We thank Erik Andersen and members of the Groen lab for helpful discussions and feedback on the manuscript. This study was funded by the US National Science Foundation (grant DGE-1922642 “NRT: Plants-3D (Discover, Design and Deploy): Enhancing Minority Graduate Training in Plant Synthetic Biology”) and the US National Institute of General Medical Sciences of the National Institutes of Health (grant R35GM151194).

---

**Figure S1:**
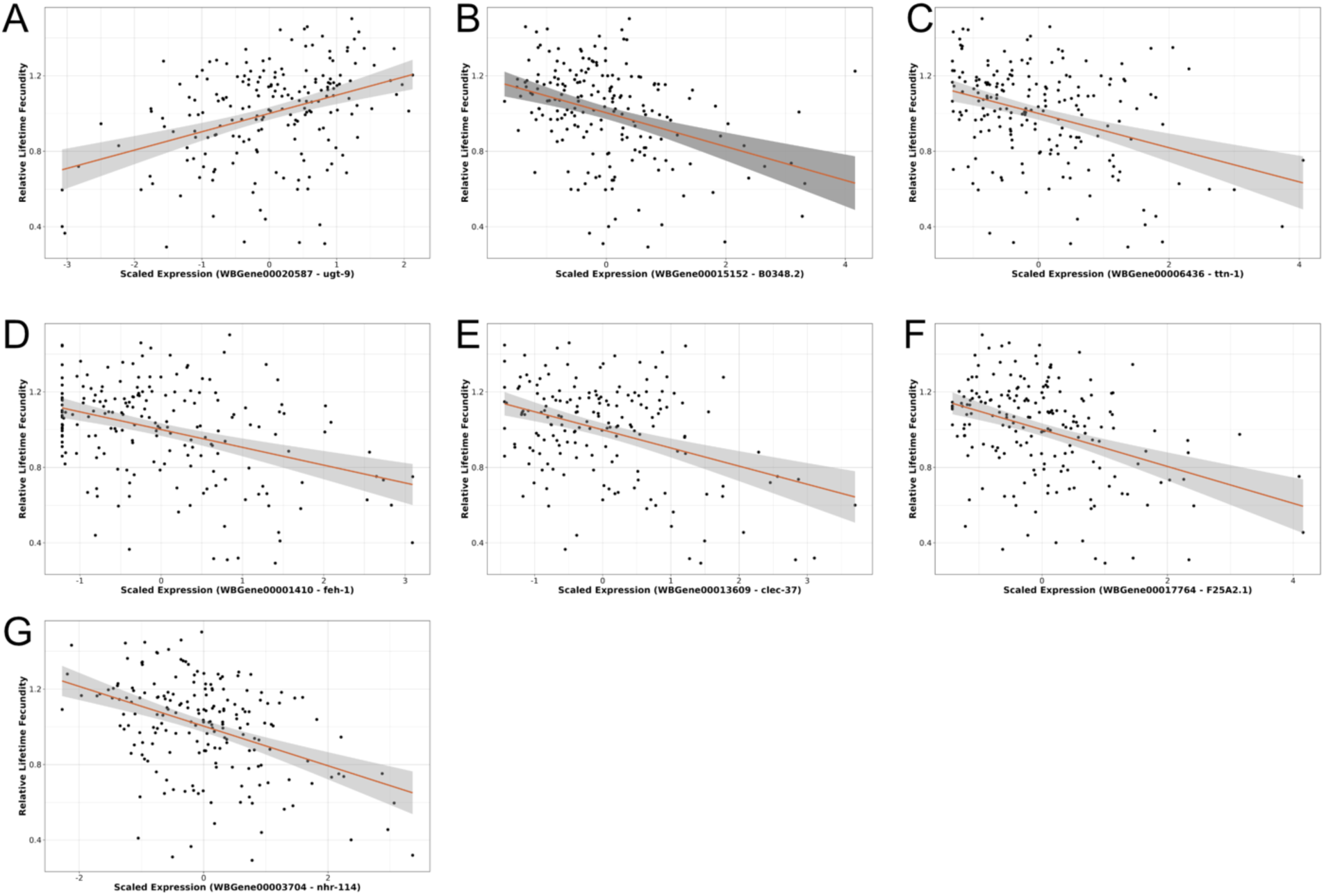
Transcripts with abundances that show significant covariance with fecundity. Regressions of fecundity fitness on abundances of seven transcripts that retained significant linear selection differentials (*S*) following Bonferroni correction (p < 3.87 x 10^-7^), including transcripts encoding **A.** a UDP-glucuronosyltransferase involved in detoxification of dietary metabolites, **B.** a predicted Zinc ion-binding protein, **C.** a major component of the muscle actin filament, **D.** the *C. elegans* homolog of the Fe65 protein implicated in Alzheimer’s disease, **E.** an uncharacterized C-type lectin, **F.** an uncharacterized protein with a predicted role in lipid metabolism, and **G.** a nuclear hormone receptor that buffers germline stem cells from oxidative damage.

**Figure S2:**
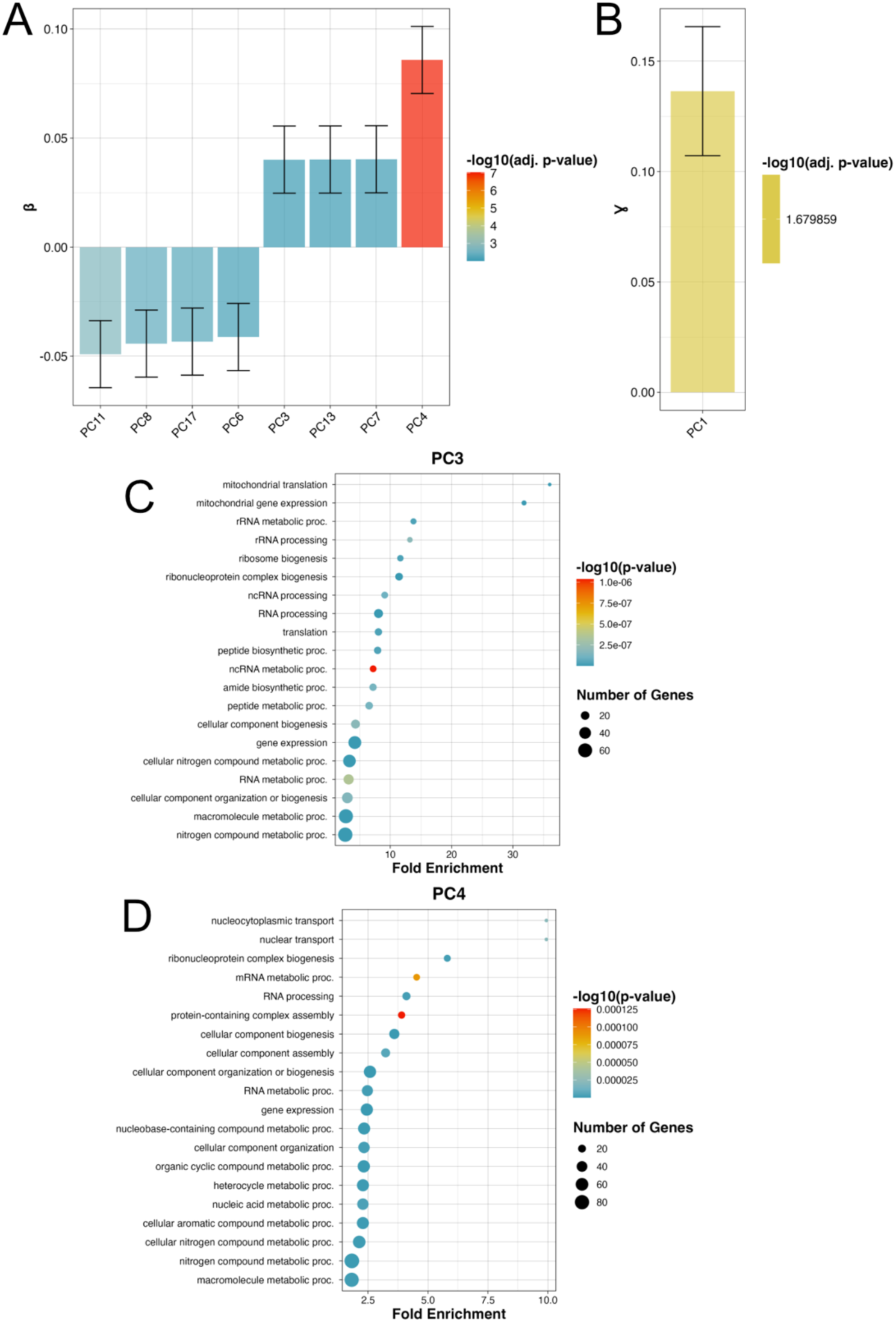
Direct effects of natural selection on correlated suites of gene expression. Multivariate selection analyses revealed that expression of suites of genes loading into **A.** eight principal components (PCs) showed significant linear selection (β) gradients (p < 0.05) and **B.** one principal component showed a significant quadratic selection (Ɣ) gradient (p < 0.05). **C, D.** Gene Ontology biological process enrichment of the top 100 genes loading most strongly into **C.** PC3 and **D.** PC4 reveals positive directional selection on correlated suites of genes involved in RNA biogenesis and post-transcriptional regulation.

**Figure S3:**
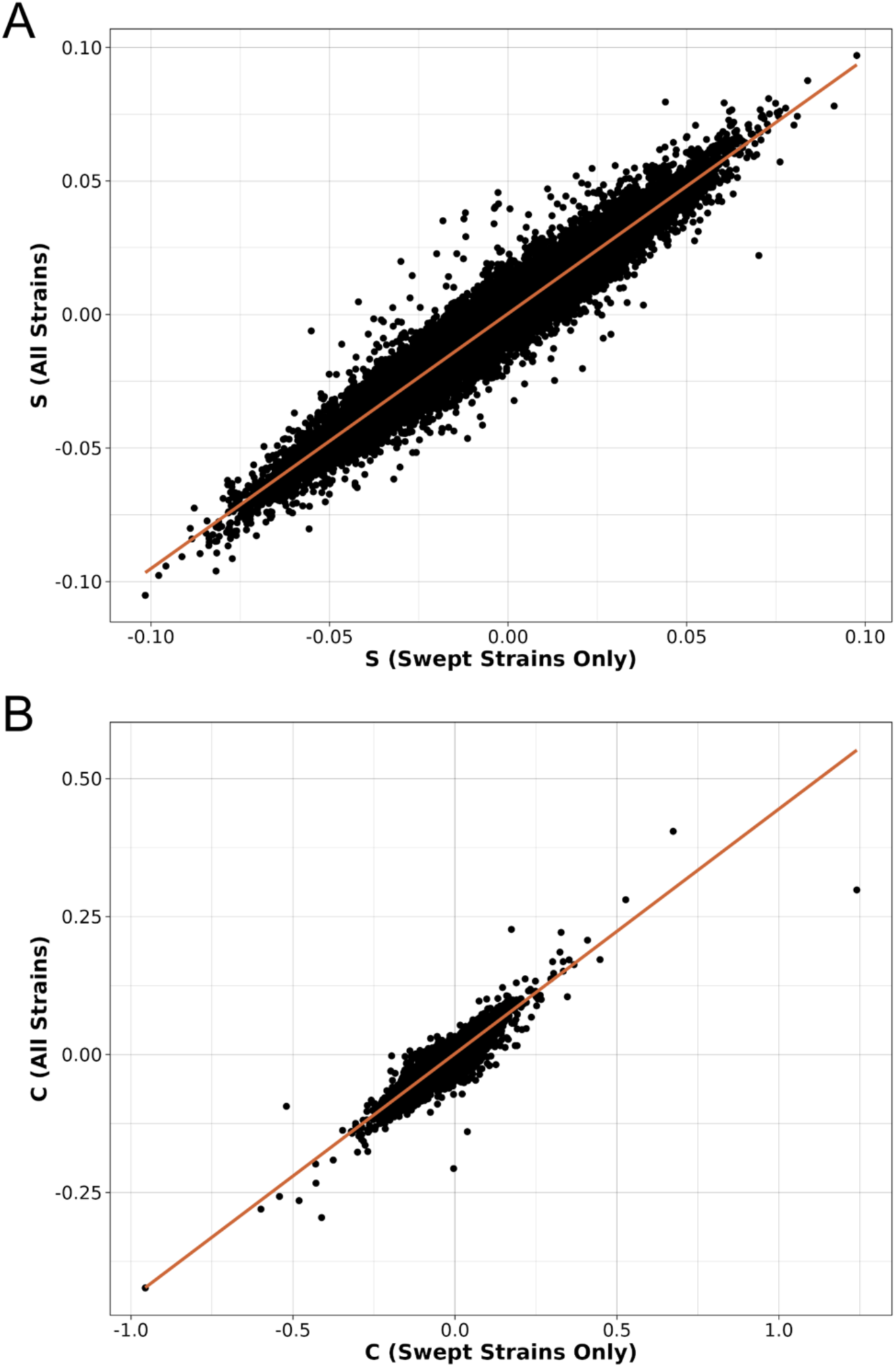
Population structure has limited impact on the observed patterns of selection. Univariate selection models were determined using only a subset of strains (n = 176) that had not experienced a historical selective sweep that caused genomic divergence of another subset of strains as determined by Zhang *et al*. (2021). Comparisons of models that included or excluded the genomically diverged strains reveal high concordance for both **A.** *S* (Spearman 𝜌 = 0.9668, p < 2.2×10^-16^) and **B.** *C* (Spearman 𝜌 = 0.9382, p < 2.2×10^-16^).

**Figure S4:**
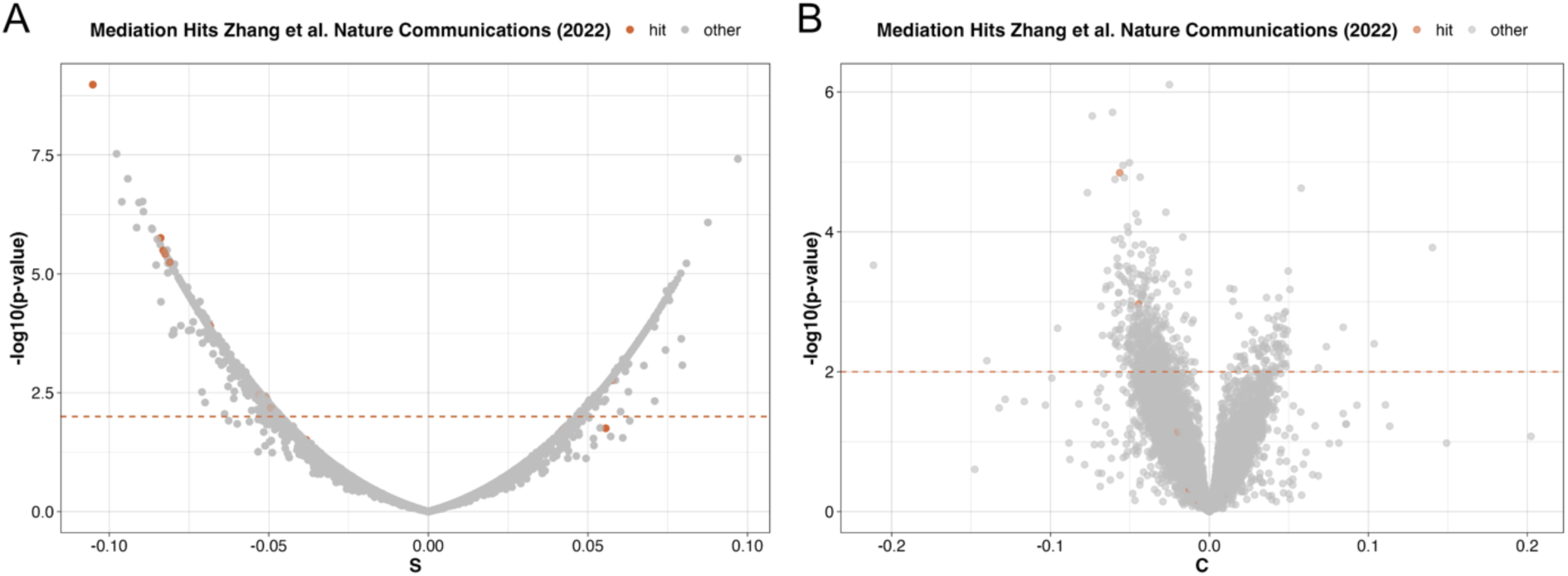
Selection models capture fecundity-associated quantitative trait loci. **A, B.** Volcano plots with genes that represent quantitative trait loci (QTL) for total lifetime fecundity, as determined by Zhang *et al*. (2022), denoted as orange-colored dots. **A.** Ten QTL encode at least one transcript with a significant linear selection differential *S*, and **B.** two QTL encode at least one transcript with a significant quadratic differential *C*. Horizontal dashed orange lines represent the threshold for statistical significance for *S* or *C*.

**Figure S5:**
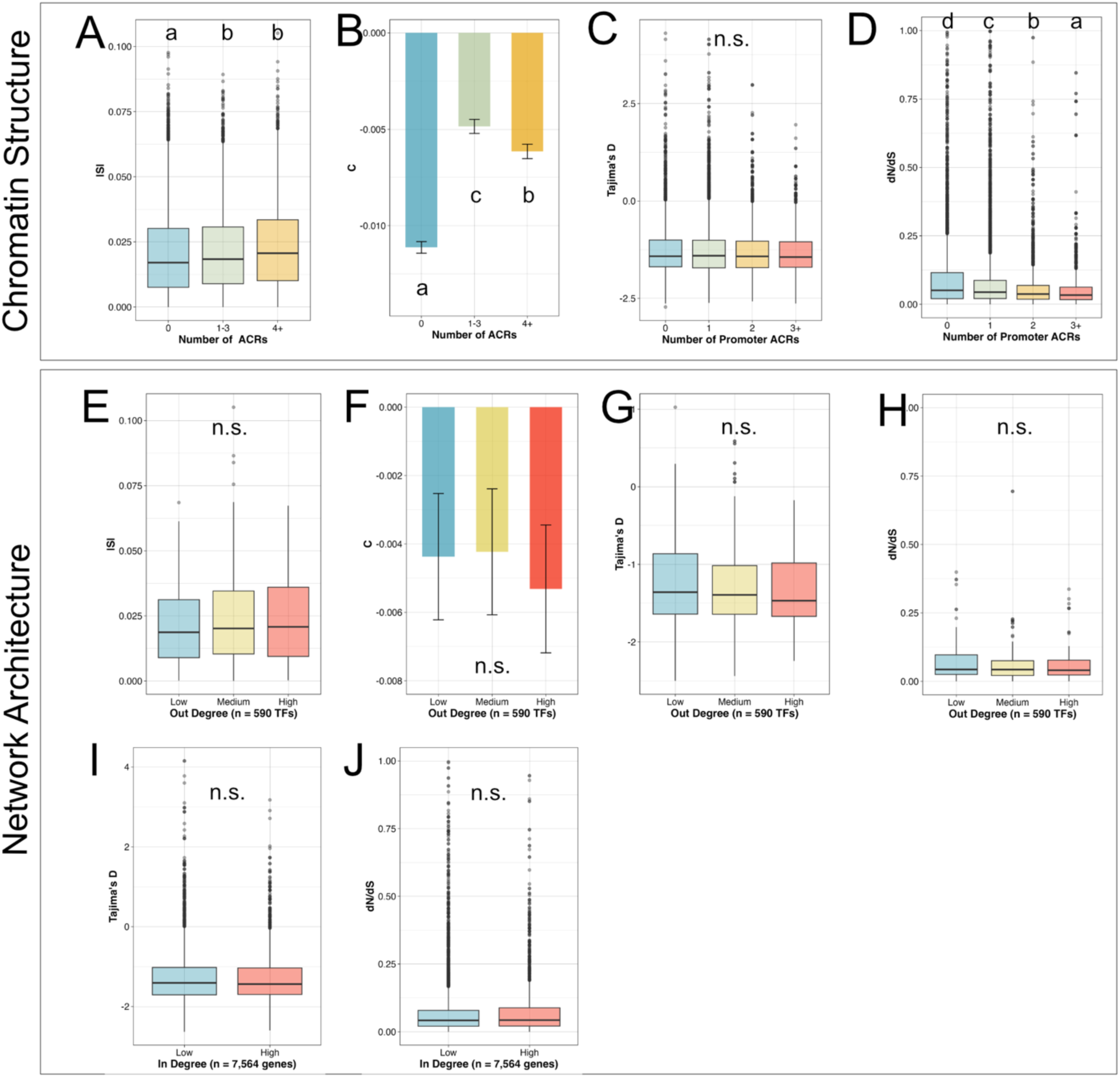
Network architecture partially uncouples patterns of sequence and expression evolution. **A, B.** Expression of genes with more accessible chromatin regions (ACRs) around enhancers experiences **A.)** stronger directional selection (Kruskal-Wallis test with Dunn’s post-hoc test, p < 2.732×10^-6^) and is further **B.)** released from constraints of stabilizing selection (Kruskal-Wallis test with Dunn’s post-hoc test, p < 0.0136). **C.** Promoter ACRs are not associated with divergent rates of genome sequence evolution (Kruskal-Wallis test, p = 0.3215), but **D.)** genes with more promoter ACRs tend to have more-slowly-evolving protein coding sequences (Kruskal-Wallis test with Dunn’s post-hoc test, p < 3.801×10^-26^). **E-H.** The number of genes transcription factors (TFs) regulate is not associated with the strength of **E.** directional (Kruskal-Wallis tests, p = 0.3756) or **F.** quadratic selection on expression (p = 0.9406), **G.** rates of purifying selection on their genomic regions (p = 0.2331) or **H.** rates of evolution of their protein coding sequences (p = 0.4346). **I, J.** Genes regulated by more TFs do not diverge from genome-wide rates of **I.)** purifying selection (Mann-Whitney *U*-tests, two-sided, p = 0.2324) or **J.** protein coding sequence evolution (p = 0.08082).

**Figure S6:**
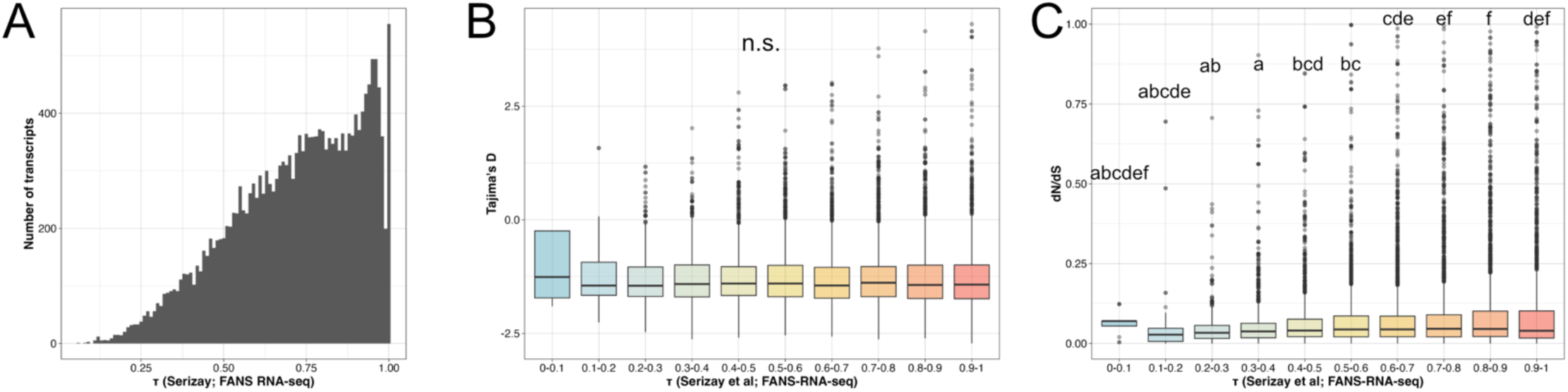
Tissue-specific genes evolve faster at the sequence level. **A.** Distribution of τ (tau) values as determined from Fluorescence-Activated Nuclei Sorting (FANS) RNA-sequencing data (Serizay *et al*. 2020). Most transcripts have tissue-enriched expression patterns. **B.** Genomic regions encoding tissue-enriched transcripts do not experience deviations from genome-wide patterns of purifying selection (Kruskal-Wallis test, p = 0.008209, none were significant after Dunn’s post-hoc tests). **C.** Genes with tissue-specific expression patterns evolve faster at the coding sequence level than those with constitutive expression patterns (Kruskal-Wallis test with Dunn’s post-hoc test, p < 2.310×10^-4^).

**Figure S7:**
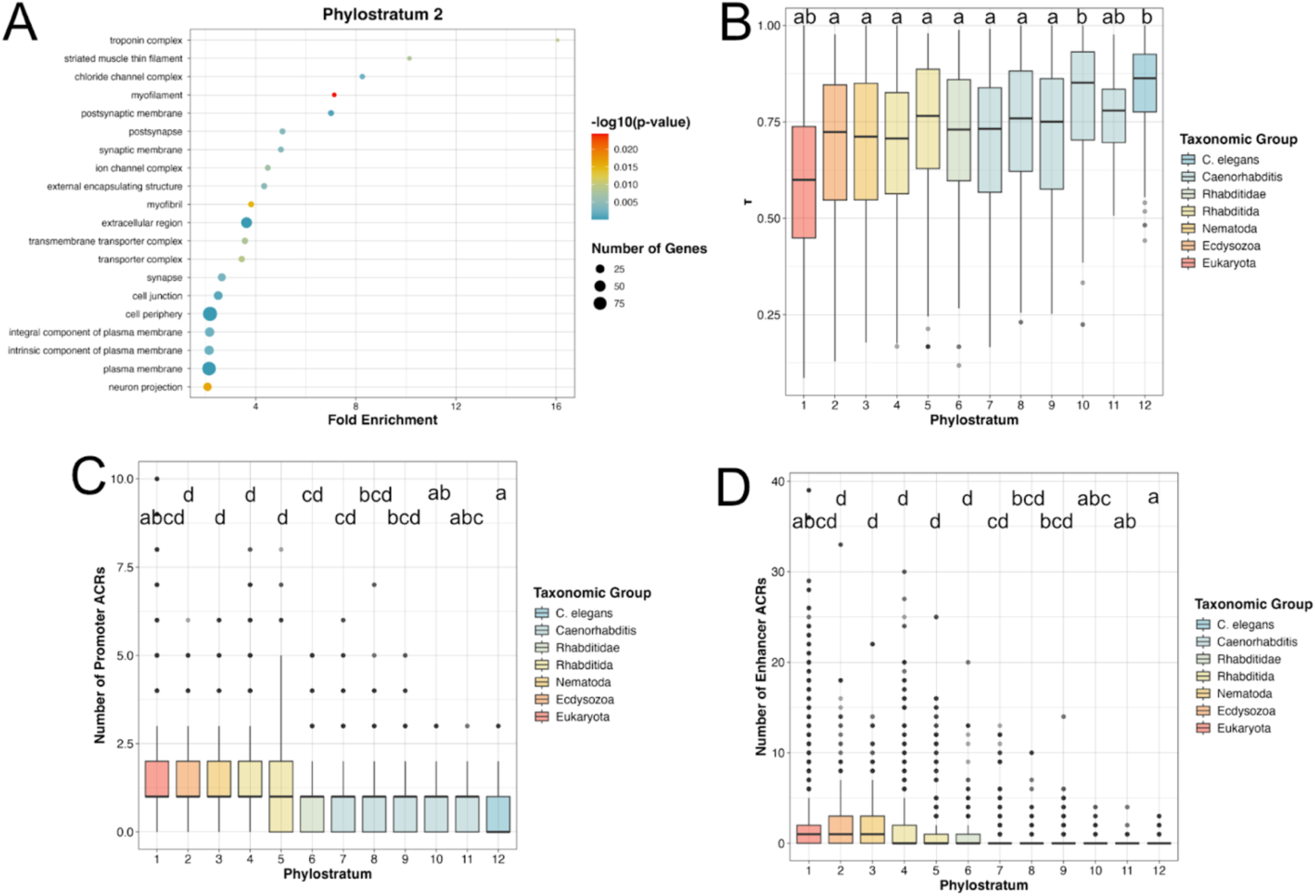
Younger genes are more likely to be tissue-specific and have less complex regulatory grammar. **A.** Genes assigned to phylostratum 2 are enriched for Gene Ontology cellular components important for the function of the neuromuscular system. **B.** Younger genes are more likely to adopt tissue-specific expression profiles (Kruskal-Wallis test with Dunn’s post-hoc test, p < 3.022×10^-4^). **C, D.** Younger genes have fewer accessible chromatin regions (ACRs) for both **C.)** promoters (Kruskal-Wallis tests with Dunn’s post-hoc tests, p < 4.685×10^-4^) and **D.)** enhancers (p < 4.693×10^-5^).

**Figure S8:**
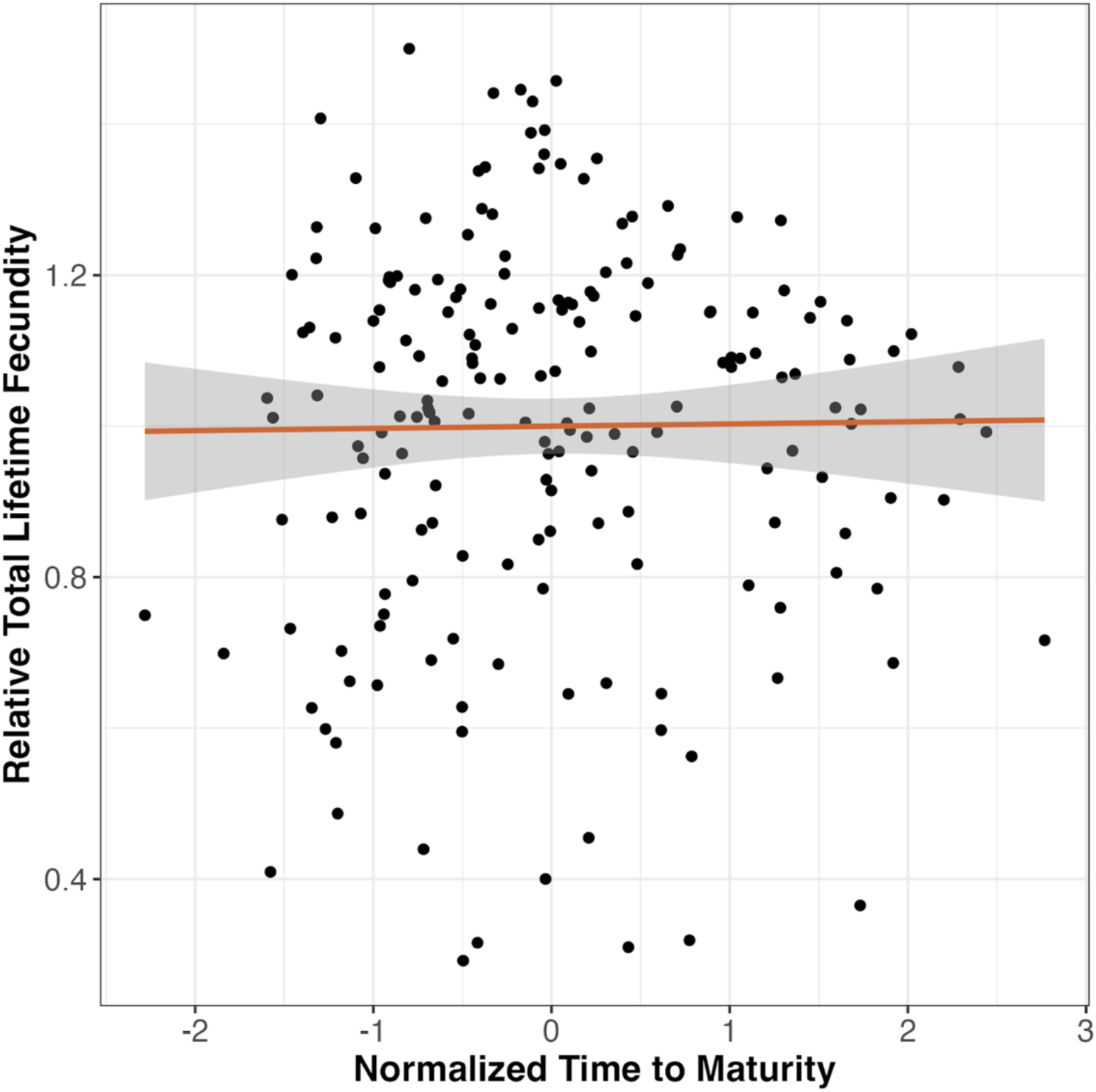
No linear covariation between maturation timing and total lifetime fecundity. Maturation timing is not under directional selection, as evidenced by the lack of a significant association with total lifetime fecundity (*S = 0.002955 ± 0.018596, p = 0.874*).

